# CAR/Nr1i3 directs T cell adaptation to bile acids in the small intestine

**DOI:** 10.1101/2020.07.30.229864

**Authors:** Mei Lan Chen, Xiangsheng Huang, Hongtao Wang, Courtney Hegner, Yujin Liu, Jinsai Shang, Amber Eliason, HaJeung Park, Blake Frey, Guohui Wang, Sarah A. Mosure, Laura A. Solt, Douglas J. Kojetin, Alex Rodriguez-Palacios, Deborah A. Schady, Casey T. Weaver, Matthew E. Pipkin, David D. Moore, Mark S. Sundrud

## Abstract

Bile acids (BAs) are lipid emulsifying metabolites synthesized in hepatocytes and maintained *in vivo* through enterohepatic circulation between the liver and small intestine^1^. As detergents, BAs can cause toxicity and inflammation in enterohepatic tissues^2^. Nuclear receptors maintain BA homeostasis in hepatocytes and enterocytes^3^, but it is unclear how mucosal immune cells tolerate high BA concentrations in the small intestine lamina propria (siLP). We previously reported that CD4^+^ T effector (Teff) cells upregulate expression of the xenobiotic transporter MDR1/*ABCB1* in the siLP to prevent BA toxicity and suppress Crohn’s disease-like small bowel inflammation^4^. Here, we identify the nuclear xenobiotic receptor, constitutive androstane receptor (CAR/*NR1I3*), as a regulator of MDR1 expression in T cells, and safeguard against BA toxicity and inflammation in the small intestine. CAR was activated and induced large-scale transcriptional reprograming in Teff cells infiltrating the siLP, but not the colon. CAR induced expression of detoxifying enzymes and transporters in siLP Teff cells, as in hepatocytes, but also the key anti-inflammatory cytokine, *Il10*. Accordingly, CAR-deficiency in T cells exacerbated, whereas pharmacologic CAR activation suppressed, BA-driven ileitis in T cell-reconstituted *Rag*^−/−^ mice. These data suggest that CAR acts locally in small intestinal T cells to detoxify BAs and resolve inflammation. Activation of this program offers an unexpected strategy to treat small bowel Crohn’s disease, and provides evidence of lymphocyte sub-specialization within the small intestine.

---

Seeking transcriptional mechanisms underlying MDR1 upregulation in siLP Teff cells^4^, we considered the ligand-regulated nuclear receptors (NRs), a family of environmental-sensing transcription factors that control diverse gene expression programs important for immunity, inflammation, metabolism and gastrointestinal physiology^5^. To assess functions of all 49 mouse NRs, their individual contributions to mucosal Teff cell MDR1 expression were assessed in a pooled *in vivo* RNAi screen. Activated naïve CD4^+^ T cells transduced separately with 258 retroviruses expressing shRNAmirs against 70 genes (Supplemental Table 1) were pooled, FACS-sorted for retroviral reporter (Ametrine) expression, and transferred into syngeneic (FVB/N, ‘FVB’) *Rag1*^−/−^ mice. Six-weeks later, transduced Teff cells were recovered from spleen or siLP, and MDR1^hi^ or MDR1^lo^ subsets were isolated based on *ex vivo* efflux of the fluorescent MDR1 transport substrate, rhodamine 123 (Rh123)^6^. shRNAmir abundances were quantified by DNA-seq (**Fig. 1a**).

**Fig. 1.**
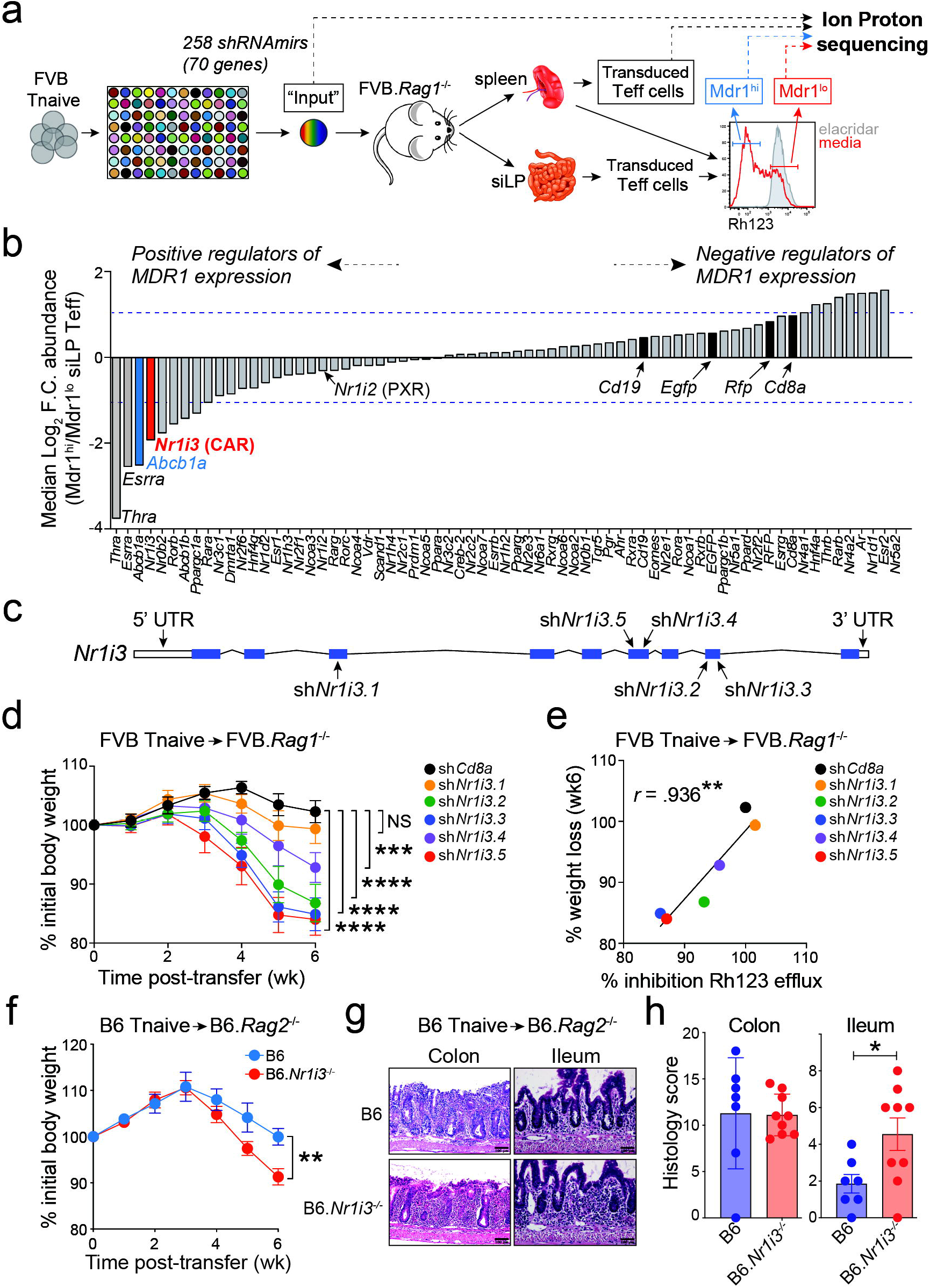
CAR/Nr1i3 regulates MDR1 expression in mucosal CD4^+^ T cells. **(a)** Schematic outline of the pooled *in vivo* RNAi screening approach (see Methods for details). **(b)** Median log_2_ fold-change in shRNAmir abundances in MDR1^hi^ *vs*. MDR1^lo^ siLP Teff cells, determined by DNA-seq and grouped by gene (see Supplemental Table 1 for the list of shRNAmirs used in the screen). Horizontal lines indicate 2-fold changes. **(c)** Diagram of the *Nr1i3*/CAR locus. Position of seed sequences for each CAR/*Nr1i3*-specific shRNAmir is shown; 5’ and 3’ untranslated regions (UTR); filled boxes depict exons. **(d)** Mean weight loss (± SEM) in co-housed FVB.*Rag1*^−/−^ mice receiving FVB wild type CD4^+^ T cells expressing negative control *shCd8a* (*n* = 11), or CAR/*Nr1i3*-specific shRNAmirs; *shNr1i3.1* (*n* = 7), *shNr1i3.2* (*n* = 7), *shNr1i3.3* (*n* = 7), *shNr1i3.4* (*n* = 7), *shNr1i3.5* (*n* = 7). \*\*\**P* < .001, \*\*\*\**P* < .0001, Two-way ANOVA. Data from two experiments. **(e)** Correlation between severity of T cell transfer-induced weight loss (at 6-weeks post-T cell transfer; as in [d]) and MDR1-dependent Rh123 efflux in *ex vivo*-isolated spleen Teff cells (determined by flow cytometry; see Extended Data Fig. 1c-d). \*\**P* < .01, Pearson correlation test. **(f)** Mean weight loss (± SEM) in co-housed B6.*Rag2*^−/−^ recipients of C57BL/6 wild type (B6; blue; *n* = 7) or CAR-deficient (B6.*Nr1i3*^−/−^; red; *n* = 9) naïve CD4^+^ T cells. \*\**P* < .01, Two-way ANOVA. Data from two experiments. **(g)** H&E-stained sections of colons or terminal ilea from transferred B6.*Rag2*^−/−^ mice (at 6-weeks post-T cell transfer; as in [f]). Representative of 7-9 mice per group analyzed over two experiments. **(h)** Mean histology scores (± SEM) for colons or terminal ilea as in (f-g). B6.*Rag2*^−/−^ mice receiving wild-type (B6; *n* =7) or CAR-deficient (B6.*Nr1i3*^−/−^; *n* = 9) T cells \**P* < .01, unpaired two-tailed student’s *t* test.

Multiple shRNAmirs against constitutive androstane receptor (CAR/*Nr1i3*) and MDR1/*Abcb1a* itself were enriched in MDR1^lo^ *vs*. MDR1^hi^ Teff cells from both spleen and siLP (**Fig. 1b**; Extended Data Fig. 1a-b). As CAR prevents BA-induced hepatotoxicity^8^, and regulates hepatic MDR1 expression^9^, these results suggested CAR might have similar protective functions in Teff cells infiltrating the siLP. We confirmed 3 of 5 CAR/*Nr1i3*-specific shRNAmirs reduced MDR1-dependent Rh123 efflux in Teff cells recovered from transferred *Rag1*^−/−^ mice (**Fig. 1c**, Extended Data Fig. 1c-d). These same clones silenced CAR/*Nr1i3* expression, as judged by *ex vivo* qPCR, and reduced expression of both MDR1/*Abcb1a* and the signature CAR target gene, *Cyp2b10*^10^ (Extended Data Fig. 1e).

CAR regulates transcription as a heterodimer with retinoid X receptors (RXRα/β/γ)^11^. However, RXRs also dimerize with other NRs that regulate diverse aspects of T cell function *in vivo*^11^. Accordingly, shRNAmir-mediated RXRα depletion predominantly impacted Teff cell persistence *in vivo* (Extended Data Fig. 1f). Depletion of the CAR-related xenobiotic-sensor, pregnane X receptor (PXR/*Nr1i2*)^12^, had little influence on either MDR1 expression or Teff cell persistence (**Fig. 1b**; Extended Data Fig. 1f). Consistent with this, Teff cells from C57BL/6 (B6)-derived CAR-deficient (*Nr1i3*^−/−^) mice, but not PXR-deficient (*Nr1i2*^−/−^) mice, displayed lower MDR1 expression than bystander CD45.1 wild type cells after co-transfer into *Rag1*^−/−^ mice; cells lacking only CAR showed equivalently low MDR1 expression as those lacking both CAR and PXR (Extended Data Fig. 1g-i). These data implicate CAR in the regulation of mucosal T cell function *in vivo*.

The degree to which shRNAmir-mediated CAR depletion attenuated MDR1 expression in FVB wild type Teff cells transplanted into *Rag1*^−/−^ mice correlated directly with the severity of weight loss these cells induced (**Fig. 1d-e**; Extended data Fig. 1c-d). This was consistent with our prior observation that FVB T cells lacking MDR1 (*Abcb1a*^−/−^*Abcb1b*^−/−^) induce more severe weight loss than wild type counterparts in reconstituted *Rag1*^−/−^ mice—due to induction of both colitis and BA-driven ileitis^4^— and is distinct from wild type naïve CD4^+^ T cells, which induce only colitis in immunodeficient hosts^13^. Naive T cells from B6-derived CAR-deficient mice also produced increased weight loss and ileitis than wild type counterparts, but equivalent colitis, after transfer into *Rag2*^−/−^ mice co-housed to normalize microflora (**Fig. 1f-h**). Therapeutic administration of cholestyramine (CME)^14^, a BA sequestering resin that prevents BA reabsorption into the siLP, normalized weight loss and ileitis between *Rag2*^−/−^ recipients of wild type or CAR-deficient T cells (Extended Data Fig. 2a-b); as did ablation of the ileal BA reuptake transporter, Apical sodium-dependent BA transporter (Asbt/*Slc10a2*)^15^, in *Rag1*^−/−^ recipients (Extended Data Fig. 2c-d). Neither genetic nor pharmacologic inhibition of ileal BA reabsorption affected severity of T cell transfer-induced colitis (Extended Data Fig. 2b, 2d). These results suggest CAR acts selectively in T cells to regulate small bowel immune homeostasis; CAR-deficiency in T cells exacerbates ileitis that is not transmissible by microbiota and requires BA reabsorption.

To elucidate CAR-dependent transcriptional programs in T cells, bystander CD45.1 wild type and CD45.2 CAR-deficient Teff cells were purified from spleen, siLP or colon lamina propria (cLP) of co-transferred *Rag1*^−/−^ mice and analyzed by RNA-seq (**Fig. 2a**). Gene expression in wild type Teff cells differed substantially between spleen, siLP and cLP, whereas CAR-deficiency most conspicuously altered gene expression in siLP Teff cells (**Fig. 2b**). CAR-deficient siLP Teff cells failed to upregulate many ‘siLP-signature’ genes preferentially expressed in wild type cells from siLP *vs*. either spleen or cLP, and ectopically expressed genes characteristic of wild type Teff cells from colon (**Fig. 2c-d**). siLP-signature genes encoding chaperones, receptors and enzymes involved in lipid binding, transport and metabolism (*e.g*., *Apold1*, *Pex26*, *Dgkh*, *Ldlr*, *Phyhd1*, *Lclat1*) were among those decreased in CAR-deficient *vs*. wild type siLP Teff cells (**Fig. 2c-d**, Supplemental Table 2); as were genes induced by CAR in mouse hepatocytes after *in vivo* administration of the specific agonist ligand, 1,4-*Bis*(3,5-Dichloro-2-pyridinyloxy) benzene (TCPOBOP, ‘TC’)^16^ (Extended Data Fig. 3a-b). Genes showing CAR-dependent expression in both siLP Teff cells and hepatocytes were enriched for loci at which TC-inducible CAR DNA-binding has been observed in hepatocytes by ChIP-seq^17^ (Extended Data Fig. 3c). These included MDR1/*Abcb1a* and *Cyp2b10*, as expected, but also other ABC-family transporters (*e.g*., *Abcb4*) and cytochrome P450 enzymes (*e.g.*, *Cyp2r1*) (Extended Data Fig. 3d), suggesting that CAR activates a ‘hepatocyte-like’ BA-detoxification program in siLP Teff cells. CAR-deficient Teff cells accumulated less in the siLP of *Rag1*^−/−^ recipients initially, and in all tissues later, relative to wild type bystanders (Extended Data Fig. 4a-c). Ablating Asbt-dependent BA reabsorption in *Rag1*^−/−^ recipients tended to minimize this phenotype (Extended Data Fig. 4d-f).

**Fig. 2.**
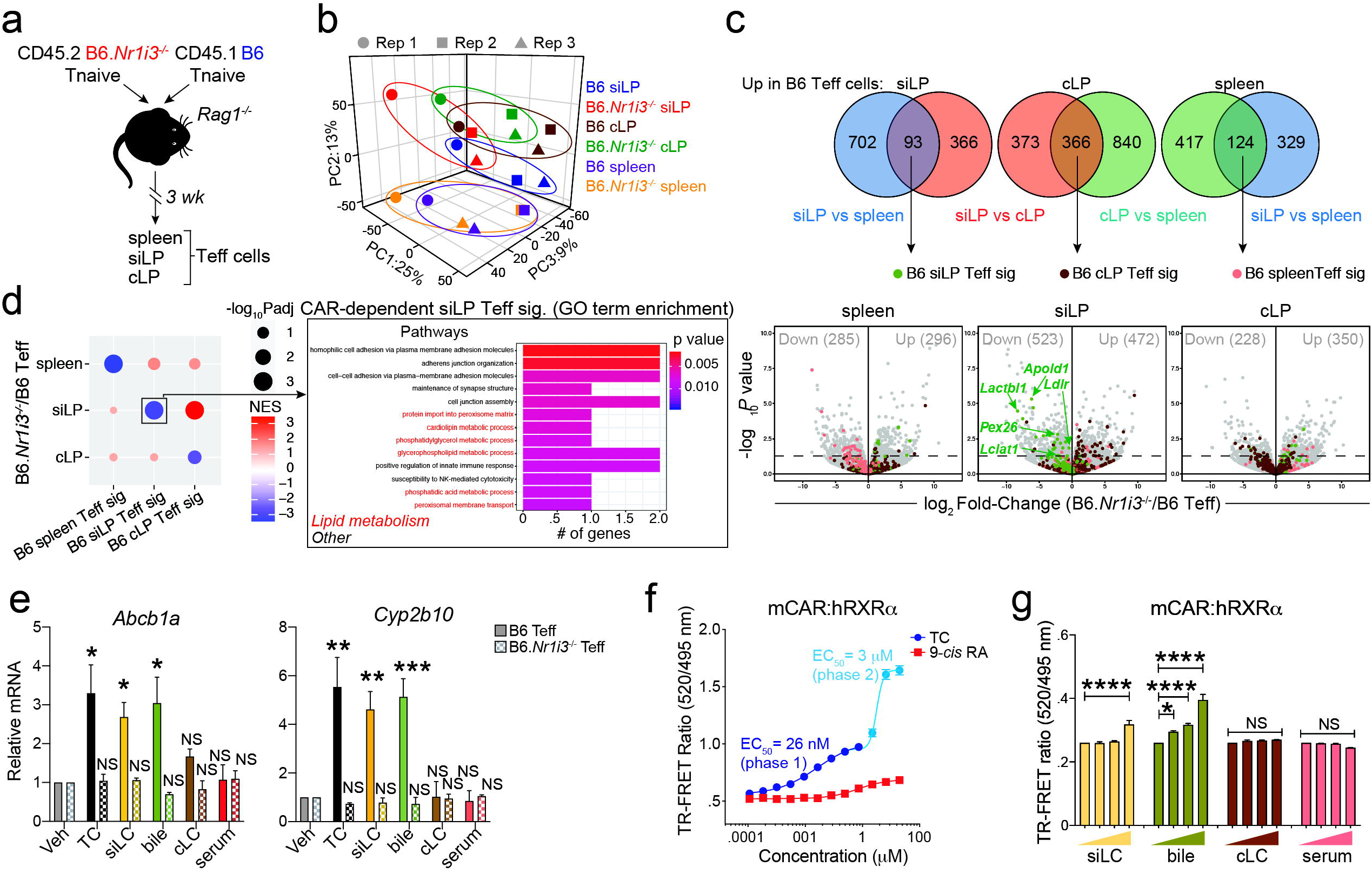
CAR regulates T cell gene expression in the small intestine. **(a)** Mixed T cell transfer approach for RNA-seq analysis of wild type (B6) and CAR-deficient (B6.*Nr1i3*^−/−^) T effector (Teff) cells. **(b)** PCA of wild type and CAR-deficient Teff cell gene expression in spleen, small intestine lamina propria (siLP) or colon lamina propria (cLP). **(c)** *Top*, identification of signature (sig) genes expressed highest in spleen, siLP or cLP wild type Teff cells. Comparisons, and numbers of genes higher (sample 1/2) in each comparison, are listed. *Bottom*, differential gene expression between spleen, siLP or cLP wild type and CAR-deficient Teff cells. Numbers of Up/Down genes are indicated. Tissue-specific Teff cell-signature genes are highlighted/annotated. **(d)** *Left*, enrichment of tissue-specific Teff cell-signature genes (x-axis; as in [c]) within those differentially expressed between wild type and CAR-deficient cells per tissue (y-axis). *Right*, Gene ontology (GO) terms enriched in CAR-dependent siLP Teff signature genes; lipid metabolic pathways highlighted red. (b-d) Gene expression data from 3-independent experiments. **(e)** Mean relative *Abcb1a* or *Cyp2b10* expression (± SEM; *n* = 3), by qPCR, in *ex vivo*-isolated wild type or CAR-deficient Teff cells stimulated +/− mouse tissue extracts. Veh, vehicle; TC, TCPOBOP (CAR agonist); siLC, small intestine lumen content; cLC, colon lumen content. \**P* < .05, \*\**P* < .01, \*\*\**P* < .001, One-way ANOVA with Dunett’s correction for multiple comparisons. NS, not significant. **(f)** Mouse (m)CAR:human (h)RXRα ligand-binding domain (LBD) heterodimer activation (± SEM; *n*=3), by TR-FRET, +/− TC or 9-*cis* retinoic acid (RA, hRXRα agonist). Numbers indicate EC_50_’s; representative of 5 experiments. **(g)** Mean mCAR:hRXRα LBD heterodimer activation (± SEM; *n* = 3), by TR-FRET, +/− mouse tissue extracts. (*L-R*): 0%; 0.01%, 0.1%,, 1% extract. Incorporates data from 3 experiments. \**P* < .05, \*\*\*\**P* < .0001, one-way ANOVA with Tukey’s correction for multiple comparisons. NS, not significant.

To test if CAR also regulates small intestine-associated T cell function in humans, we analyzed CAR expression and function in healthy adult peripheral blood T cell subsets most likely to have recirculated from the siLP. 1-5% of circulating Teff cells expressed the requisite combination of receptors for siLP-homing, α4β7 integrin and CCR9^18^ (α4^+^β7^+^CCR9^+^), whereas naïve T cells that lack gut-homing potential did not (Extended Data Fig. 5a-c). Fewer CD25^+^ T regulatory (Treg) cells expressed these receptors (Extended Data Fig. 5a-c), suggesting Treg cells may be more efficiently retained in the siLP than Teff cells. As predicted, siLP-linked α4^+^β7^+^CCR9^+^ Teff cells displayed elevated expression of MDR1/*ABCB1*, CAR/*NR1I3* and *CYP2B6* (ortholog of mouse *Cyp2b10*^19^), compared with naïve, Treg or Teff cells lacking one or more siLP-homing receptors (Extended Data Fig. 5d-f). In addition, only α4^+^β7^+^CCR9^+^ Teff cells responded to *ex vivo* treatment with the human CAR agonist ligand, 6-(4-Chlorophenyl)imidazo[2,1-b][1,3]thiazole-5-carbaldehyde O-(3,4-dichlorobenzyl) oxime (CITCO)^19^ by upregulating *CYP2B6* and *ABCB1* (Extended Data Fig. 5g-h). CCR6^+^CXCR3^hi^CCR4^lo^ “Th17.1” cells—which possess both Th17 and Th1 effector functions, and high MDR1 expression^20^—were enriched among total α4^+^β7^+^CCR9^+^ Teff cells (Extended Data Fig. 5i-l). However, α4^+^β7^+^CCR9^+^ Th17.1 cells exhibited higher MDR1 expression than Th17.1 cells lacking one or more siLP-homing receptors (Extended Data Fig. 5m-n). The same was true for Th17 (CCR6^+^CXCR3^lo^CCR4^hi^) and Th1 (CCR6^−^CXCR3^hi^ CCR4^lo^) cells. These data suggest that CAR preferentially operates in both mouse and human Teff cells in the small intestine.

Preferential CAR function in siLP Teff cells could involve local activation by endogenous metabolites. Consistent with this possibility, gallbladder bile or sterile-filtered soluble small intestine lumen content (siLC) from wild type B6 mice—but not colon lumen content (cLC) or serum—induced MDR1/*Abcb1a* and *Cyp2b10* upregulation in *ex vivo*-stimulated wild type, but not CAR-deficient, Teff cells from transferred *Rag1*^−/−^ mice (**Fig. 2e**; Extended Data Fig. 6a). CAR-dependent gene expression in this *ex vivo* culture system was also induced by TC, inhibited by the CAR inverse agonist, 5α-Androstan-3β-ol^8^, and unaffected by the PXR agonist, 5-Pregnen-3β-ol-20-one-16α-carbonitrile (PCN)^12^ (**Fig. 2e**; Extended Data Fig. 6b). Bile and siLC concentrations that enhanced CAR-dependent gene expression in Teff cells also promoted recruitment of a PGC1α co-activator peptide to recombinant CAR-RXRα ligand-binding domain (LBD) heterodimers, but not to RXRα LBD homodimers, in time-resolved fluorescence resonance energy transfer (TR-FRET) experiments (**Fig. 2f-g**; Extended Data Fig. 7a-b). As CAR is thought to indirectly sense, but not directly bind major BA species^21^, we reasoned that biliary metabolites other than BAs might activate the CAR LBD; bile is comprised of mixed micelles containing BAs, phospholipids, cholesterol, fatty acids and bile pigments (*e.g.*, bilirubin)^1^. Indeed, siLC pre-treated with CME to deplete free BAs^4^ retained capacity to activate CAR-RXRα LBD heterodimers (Extended Data Fig. 7c). Further, no major BA species activated CAR-RXRα LBD heterodimers in TR-FRET experiments, or stimulated CAR-dependent gene expression in *ex vivo*-cultured Teff cells (Extended Data Fig. 7d, data not shown). siLC from germ-free mice also activated CAR-RXRα LBD heterodimers (Extended Data Fig. 7c, data not shown), together suggesting that host-derived, non-BA constituents of bile may stimulate local CAR transcriptional activity in siLP Teff cells.

To further explore CAR immunoregulatory functions, we examined its control of gene expression associated with major pro- and anti-inflammatory T helper cell lineages. Genes expressed selectively in type 1 regulatory (Tr1) cells^22^—a Foxp3^−^IL-10^+^ subset known for suppressing mucosal inflammation in humans and mice^23^—were enriched among those showing reduced expression in CAR-deficient *vs*. wild type siLP Teff cells (**Fig. 3a-c**). Conversely, genes characteristic of pro-inflammatory IL-17-secreting (Th17) cells^24^ were positively enriched within siLP Teff cells lacking CAR (**Fig. 3b**). In line with these signatures, CAR-deficient Teff cells inefficiently expressed both a *Thy1.1*-expressing *Il10* reporter (‘10BiT’^25^; **Fig. 3d-e**), and endogenous IL-10 protein (Extended Data Fig. 8a-e), after transfer into *Rag1*^−/−^ mice. Reduced *Il10* expression in Teff cells lacking CAR paralleled their accumulation as RORγt^+^IL-17A^−^ ‘poised’ Th17 cells^26,27^ in siLP (Extended Data Fig. 8f-g). However, *Il10*^−/−^ T cells replete for CAR recapitulated this phenotype (Extended Data Fig. 8h-i), suggesting that CAR may reciprocally regulate Tr1 and Th17 cell development in the siLP via IL-10 induction. TC (synthetic CAR agonist), as well as bile and siLC from wild type mice, each promoted *Il10* upregulation in *ex vivo*-stimulated wild type, but not CAR-deficient, Teff cells (**Fig. 3f**; Extended Data Fig. 6), akin to *Abcb1a* and *Cyp2b10* (**Fig. 2e**).

**Fig. 3.**
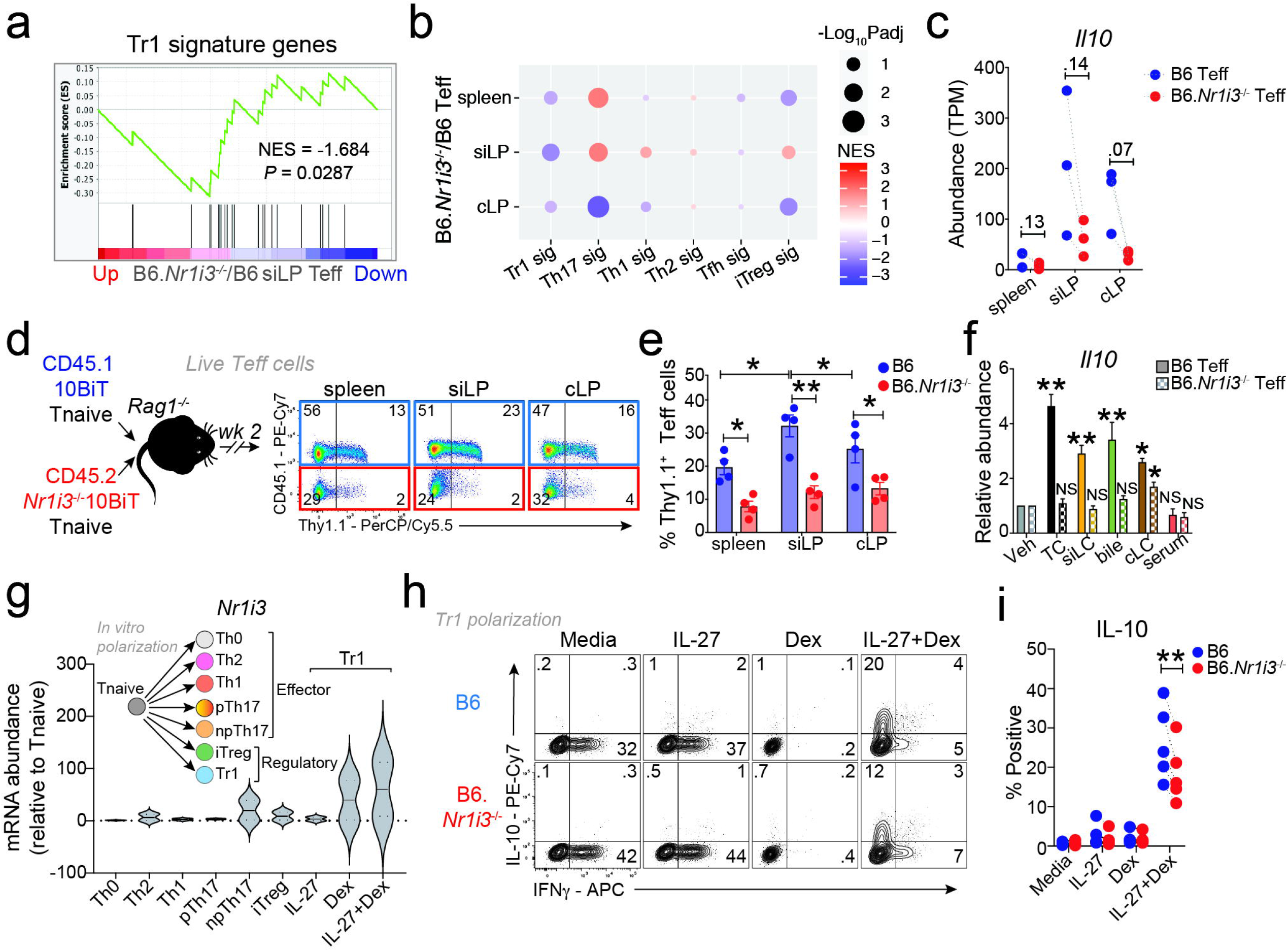
CAR promotes T cell IL-10 expression. **(a)** Type 1 regulatory (Tr1) cell-signature enrichment among genes reduced in CAR-deficient (B6.*Nr1i3*^−/−^) *vs*. wild type (B6) small intestine lamina propria (siLP) T effector (Teff) cells. NES, normalized enrichment score. **(b)** Enrichment of T cell lineage-specific genes (x-axis; see methods) within genes differentially expressed between wild type and CAR-deficient Teff cells per tissue (y-axis). **(c)** *Il10* expression, by RNA-seq (*n* = 3; see Fig. 2a), in wild type or CAR-deficient spleen, siLP, or cLP Teff cells. Paired two-tailed student’s *t* test *P* values are shown. **(d)** *Left*, mixed T cell transfer approach to analyze CAR-dependent *Il10*-reporter (Thy1.1^24^) expression in Teff cells. *Right*, *ex vivo* Thy1.1 (*Il10*) staining, in wild type or CAR-deficient Teff cells as above. Representative of 4 mice in 2-independent experiments. **(e)** Mean Thy1.1 (*Il10*)-expressing wild type or CAR-deficient Teff cell percentages (*n* = 4; ± SEM) as in (d). \**P* < .05, \*\**P* < .01, One-way ANOVA with Tukey’s correction for multiple comparisons. **(f)** Mean relative *Il10* expression (± SEM; *n* = 3), by qPCR, in *ex vivo*-isolated wild type or CAR-deficient Teff cells stimulated +/− mouse tissue extracts. Veh, vehicle; TC, TCPOBOP (CAR agonist); siLC, small intestine lumen content; cLC, colon lumen content. \**P* < .05, \*\**P* < .01, One-way ANOVA with Dunett’s correction for multiple comparisons. NS, not significant. **(g)** Box and violin plot of CAR/*Nr1i3* expression relative to CD4^+^ naïve T cells (Tnaive; *n* = 2), by qPCR, in *in vitro*-polarized wild type effector or regulatory subsets. npTh17, non-pathogenic Th17 cells; pTh17, pathogenic Th17 cells. **(h)** Intracellular IL-10/IFNγ staining in wild type or CAR-deficient Tr1-like cells. Representative of 5 experiments; numbers indicate percentages. **(i)** Mean percentages (*n* = 5; ± SEM) of IL-10^+^ Tr1-like cells as in (h). \*\**P* < .01, paired two-tailed student’s *t* test.

CAR-dependent IL-10 expression in T cell-reconstituted *Rag1*^−/−^ mice was transient—peaking 2-weeks after donor T cell engraftment and waning thereafter—and followed the kinetics of both Teff cell siLP infiltration and *ex vivo* CAR (*Nr1i3*), MDR1 (*Abcb1a*) and *Cyp2b10* gene expression (Extended Data Fig. 9a-c). This suggested that CAR expression and function in Teff cells is increased in response to inflammation, analogous to Tr1 cell dynamics *in vivo*^28^. Using an orthologous approach to induce intestinal inflammation in wild type or CAR-deficient mice—soluble anti-CD3 injection^23^— we confirmed that CAR was required for anti-CD3 (*i.e*., inflammation)-induced IL-10 upregulation by endogenous effector and regulatory T cell subsets in the siLP, but not the spleen, and was dispensable for steady-state IL-10 expression in T cells from unmanipulated mice (Extended data Figure 9d-f).

To establish a model of CAR-dependent Tr1 cell function *in vitro*, we tested CAR expression and function in naïve CD4^+^ T cells activated and expanded in culture conditions previously reported to induce differentiation of Foxp3^−^IL-10^+^ ‘Tr1-like’ cells. Combining IL-27—a cytokine that promotes Stat3-dependent IL-10 expression^22^—with the synthetic corticosteroid, dexamethasone (Dex)^29^, strongly induced CAR/*Nr1i3* and IL-10 expression, but not Foxp3 expression, in activated T cells (**Fig. 3g-I**, data not shown). Loss of CAR impaired IL-10 production by IL-27+Dex-elicited Tr1-like cells (**Fig. 3h-i**). By contrast, CAR/*Nr1i3* expression remained low during *in vitro* development of other effector (*e.g*., Th1, Th2, Th17) or Foxp3^+^ induced (i)Treg lineages, and CAR ablation had little impact on the development or function of these cells (**Fig. 3g**, Supplemental Table 3). Together, these results suggest CAR is essential for *Il10* gene regulation in Tr1 cells, which may synergize with CAR-dependent BA-detoxification to enforce small bowel immune homeostasis.

Finally, we reasoned that if CAR-deficiency in Teff cells exacerbates BA-driven small bowel inflammation, pharmacologic CAR activation might be protective. A single administration of the CAR agonist, TC, to *Rag1*^−/−^ mice reconstituted with a mixture of CD45.1 wild type and CD45.2 CAR-deficient T cells induced *Abcb1a*, *Cyp2b10* and *Il10* upregulation in wild type, but not CAR-deficient, Teff cells within 72 hr (**Fig. 4a**). Weekly TC administration reduced ileitis, but not colitis, in *Rag2*^−/−^ mice reconstituted with only wild type T cells and fed a standard 0.2% cholic acid (CA)-supplemented diet to increase the circulating BA pool and promote small bowel injury^8,12^ (**Fig. 4b-c**). CA-feeding increased morbidity in *Rag2*^−/−^ mice receiving wild type T cells, but had no obvious effects on *Rag2*^−/−^ mice in the absence of T cell transfer (**Fig. 4b**). Therapeutic effects of TC were abolished in CA-fed *Rag2*^−/−^ mice reconstituted with CAR-deficient T cells (Extended Data Fig. 10), together suggesting that BA-supplementation promotes, whereas CAR activation in T cells suppresses, experimental ileitis.

Enterohepatic circulation establishes a marked concentration gradient of BAs in the small intestine (millimolar) and colon (micromolar), which opposes that of bacteria and bacterial metabolites^1^. While antigens from enteric flora prime both pro- and anti-inflammatory T cell responses across the intestinal tract, the specific requirement for CAR-function in siLP Teff cells—defined here in an *in vivo* screen, and relative to other NRs with known regulatory functions in the colon (*e.g*., vitamin D receptor, ‘VDR’)^30^—suggests important distinctions between the immunoregulatory microenvironments of the small and large intestines. Opposing concentration gradients of bile and bacteria in the small and large intestines could be sensed by distinct sets of NRs in mucosal lymphocytes, and instruct compartmentalized regulatory functions. Microbe-induced Foxp3^+^ Treg cell development and function, for example, is most prominent in the colon and involves VDR^30^. By contrast, we show here that the BA- and xenobiotic-sensing nuclear receptor, CAR/Nr1i3, redirects gene expression in Foxp3^−^ Teff cells infiltrating the small intestine, but not the colon, to counter BA-induced toxicity and inflammation (**Fig. 4d**). Pharmacologic CAR activation could offer a new, more targeted, approach for treating small bowel Crohn’s disease, while also providing insight into lymphocyte specialization across the intestinal tract.

**Fig. 4.**
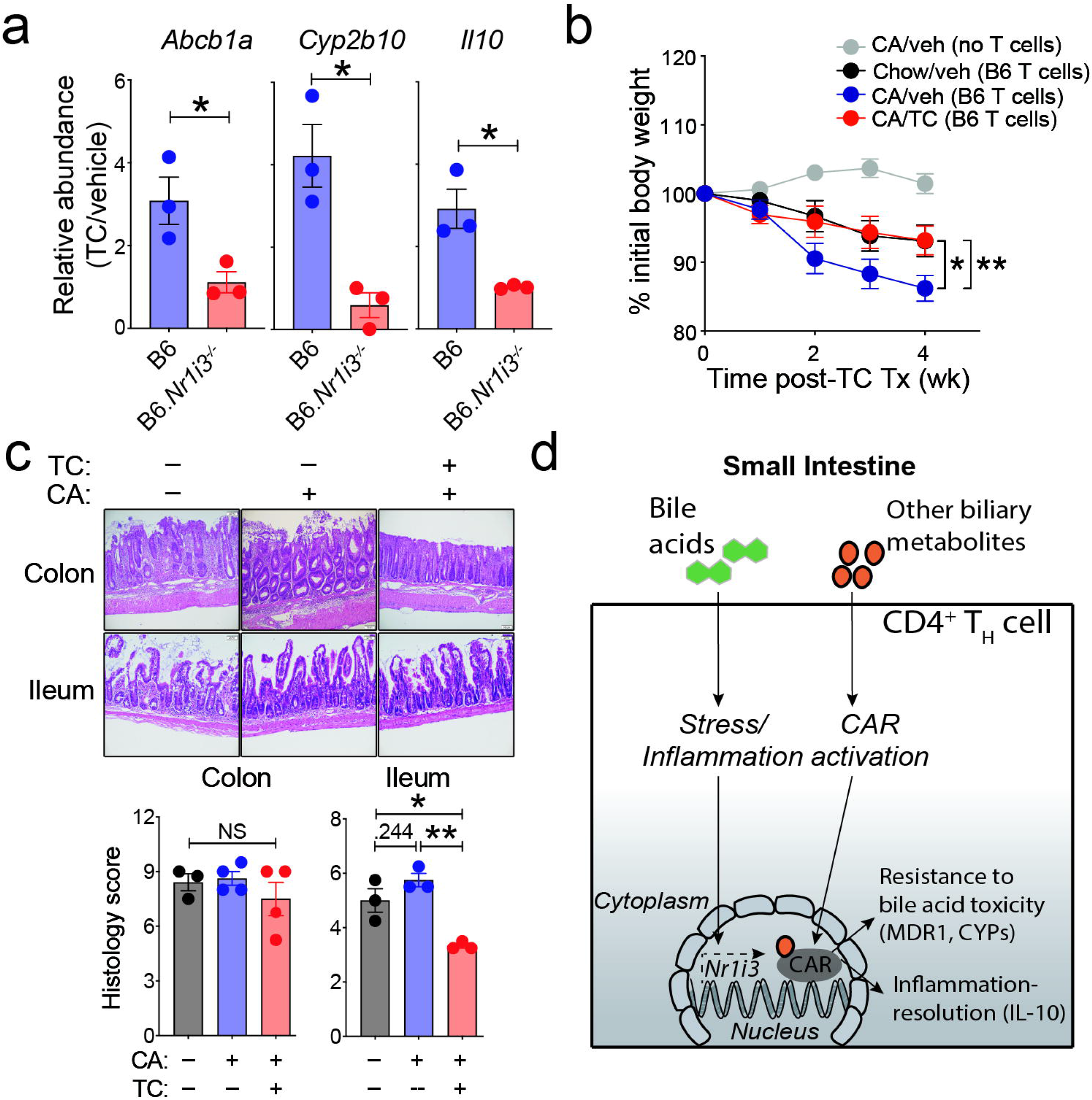
CAR activation in T cells suppresses bile acid-driven ileitis. **(a)** Mean relative *Abcb1a*, *Cyp2b10* or *Il10* expression (± SEM; *n* = 3), by qPCR, in wild type (B6) or CAR-deficient (B6.*Nr1i3*^−/−^) T effector (Teff) cells from spleens of co-transferred B6.*Rag1*^−/−^ mice 72 hr after TC (TCPOBOP; CAR agonist) or vehicle treatment. Expression in cells from TC-treated mice is shown relative to that from vehicle-treated animals; data points are from 3-independent experiments. \**P* < .05, paired two-tail student’s *t* test. **(b)** Mean weight loss (± SEM) in co-housed *Rag2*^−/−^ mice receiving wild-type naïve T cells and maintained on a CA-supplemented diet with (red; *n* = 18) or without (blue; *n* = 16) weekly TC treatment. CA-fed *Rag2*^−/−^ mice without T cell transfer (no T cells; grey; *n* =10), and *Rag2*^−/−^ mice receiving wild type T cells but left on control chow diet and treated with vehicle (black, n = 17) are also shown. Weights are relative to the start of TC treatment (3-weeks post-T cell transfer). \**P* < .05, \*\**P* < .01, Two-way ANOVA. **(c)** *Top,* H&E-stained colons or terminal ilea from *Rag2*^−/−^ mice receiving wild type T cells and fed and treated as in (b). Analyzed at week-4 post-TC treatment; representative of 3-4 mice/group. *Bottom,* mean histology scores (± SEM) for colons (*n* = 3-4) or terminal ilea (*n* = 3) as in (c). \**P* < .05, one-way ANOVA with Tukey’s correction for multiple comparisons. NS, not significant. **(d)** Model of CAR-dependent T cell regulation in the small intestine. CYPs, cytochrome P450 enzymes.

## Abbreviations

MDR1: multidrug resistance 1
IFNγ: interferon gamma
IL-10: interleukin-10
IL-17: interleukin-17
*Abcb1a*: ATP-binding cassette subfamily B, member 1a
*Abcb1b*: ATP-binding cassette subfamily B, member 1b
*Rag1/2*: recombination-activating gene 1 or 2

## Methods

### Mice

C57BL/6 (B6)-derived wild type (Stock No: 000664), CD45.1 (Stock No: 002014), *Rag1*^−/−^ (Stock No: 002216), *Rag2*^−/−^ (Stock No: 008449) and *Il10*^−/−^ (Stock No: 002251) mice were purchased from The Jackson Laboratory. Wild type FVB/N mice were purchased from Taconic. B6-derived *Nr1i2*^−/−^, *Nr1i3*^−/−^ and *Nri12*^−/−^*Nr1i3*^−/−^ mice were provided by D. Moore (Baylor College of Medicine, BCM). FVB-derived *Rag1*^−/−^ mice were a gift of Dr. Allan Bieber (Mayo Clinic, Rochester, MN). B6-derived BAC *Il10*-Thy1.1 transgenic reporter (10BiT) mice were provided by C. Weaver (University of Alabama-Birmingham, UAB) and have been described previously^24^. B6-derived *Rag1*^−/−^ mice were crossed with *Slc10a2*^−/−^ mice (gift of Dr. Paul Dawson, Emory University) in the Sundrud lab to generate *Rag1*^−/−^ mice lacking the Asbt transporter as in^4^. Lumen contents (colon, small intestine) were harvested (see below) from specific pathogen-free (SPF) or germ-free wild type B6 mice housed at the University of Alabama-Birmingham (UAB; courtesy of Dr. Weaver). All breeding and experimental use of animals was conducted in accordance with protocols approved by IACUC committees at Scripps Florida, BCM or UAB.

### Human blood samples

Human blood samples were collected and analyzed in accordance with protocols approved by Institutional Review Boards at Scripps Florida and OneBlood (Orlando, Florida). Blood was obtained following informed written consent, and consenting volunteers willingly shared clinical history and demographic information prior to phlebotomy. Institutional Review Boards at OneBlood and Scripps Florida approved all procedures and forms used in obtaining informed consent, and all documentation for consenting volunteers is stored at OneBlood.

### CD4^+^ T cell isolation and culture

Purified CD4^+^CD25^−^ T cells were magnetically isolated from spleen and peripheral lymph node mononuclear cells using an EasySep magnetic T cell negative isolation kit (Stem Cell Technologies, Inc.) with addition of a biotin anti-mouse CD25 antibody (0.5 μg/mL; BioLegend). Magnetically-enriched CD4^+^ T cells were cultured in (DMEM) supplemented with 10% heat-inactivated fetal bovine serum (BioFluids), 2mM L-glutamine (Gibco), 50uM 2-mercaptoethanol (Amresco), 1% MEM vitamin solution (Gibco), 1% MEM non-essential amino acids solution (Gibco), 1% Sodium Pyruvate(Gibco), 1% Arg/Asp/Folic acid (Gibco), 1% HEPES (Gibco), 0.1% gentamicin (Gibco) and 100u/ml Pen-Strep (Gibco). For *Rag1*^−/−^ transfer experiments, magnetically enriched CD4^+^CD25^−^ T cells were FACS-sorted to obtain pure naïve T cells (CD3^+^CD4^+^CD25^−^CD62L^hi^CD44^lo^). For *ex vivo* isolation of mononuclear cells from tissues of T cell-reconstituted Rag-deficient mice, single cell suspensions were prepared from spleen, peripheral lymph nodes, or mesenteric lymph nodes (MLN) by mechanical disruption passing through 70 μm nylon filters (BD Biosciences). For intestinal tissues, small intestines and colons were removed, rinsed thoroughly with PBS to remove the fecal contents, and opened longitudinally. Tissues were incubated for 30 minutes at room temperature in DMEM media without phenol red (Genesee Scientific) plus 0.15% DTT (Sigma-Aldrich) to eliminate mucus layer. After washing with media, intestines were incubated for 30 minutes at room temperature in media containing 1 mM EDTA (Amresco) to remove the epithelium. Intestinal tissue was digested in media containing 0.25 mg/mL liberase TL (Roche) and 10 U/mL RNase-free DNaseI (Roche) for 15-35 minutes at 37 °C. Lymphocyte fractions were obtained by 70/30% Percoll density gradient centrifugation (Sigma-Aldrich). Mononuclear cells were washed in complete T cell media and resuspended for downstream FACS analysis or sorting.

#### Naïve CD4^+^ T cell activation and polarization

magnetically enriched CD4^+^CD25^−^ T cells were seeded (at 4×10^5^ cells/cm^2^ and 1×10^6^ cells/mL) in 96- or 24-well flat bottom plates pre-coated for 2 hr at 37 °C with goat-anti-hamster whole IgG (50 μg/mL; Invitrogen). Activation was induced by adding hamster-anti-mouse CD3ε (0.3 or 1 μg/mL; BioLegend) and hamster-anti-mouse CD28 (0.25 or 0.5 μg/mL; BioXcell). After 48 hr, cells were removed from coated wells and re-cultured at 1×10^6^ cells/mL in media with or without 10 U/mL recombinant human IL-2 (rhIL-2) (NIH Biorepository), depending on the experiment (see below). For polarization studies, cells were activated in the presence of the following sets of cytokines and/or neutralizing antibodies (all from R&D Systems): Th0—media alone; Th1—recombinant human (rh)IL-12 (5 ng/mL) plus anti-mouse IL-4 (5 μg/mL); Th2—rhIL-4 (10 ng/mL) plus anti-mouse IFNγ (5 μg/mL); non-pathogenic (np)Th17—recombinant mouse (rm)IL-6 (40 ng/mL) plus rhTGFβ1 (1 ng/mL), anti-mouse IFNγ (5 μg/mL) and anti-mouse IL-4 (5 μg/mL); pathogenic (p)Th17— rmIL-6 (40 ng/mL) plus rhTGFβ1 (1 ng/mL), rhIL-23 (10 ng/mL) anti-mouse IFNγ (5 μg/mL) and anti-mouse IL-4 (5 μg/mL); induced T regulatory (i)Treg—rhTGFβ1 (5 ng/mL) plus rhIL-2 (10 U/mL), anti-mouse IFNγ (5 μg/mL) and anti-mouse IL-4 (5 μg/mL). For Tr1 cultures, cells were activated in the presence of rhIL-27 (100 ng/mL) and/or dexamethasone (100 nM; Sigma-Aldrich). Cytokine, antibodies and/or Dex were added at the time of activation (day 0), and re-added to expansion media between days 2-4 of culture. Cells were analyzed for intracellular expression of transcription factors and/or cytokines, to confirm polarization, on day 4 after re-stimulation with phorbol 12-myrisate 13-acetate (PMA; 10nM; Life Technologies) and ionomycin (1uM; Sigma-Aldrich) for 3-4 hr in the presence of brefeldin A (BFA; 10ug/mL; Life Technologies).

#### Ex vivo-stimulation of FACS-sorted effector/memory (Teff) cells from reconstituted Rag1^−/−^ mice

30,000 CD45.1 (wild type) or CD45.2 (*Nr1i3*^−/−^) cells—FACS-purified from spleens of B6.*Rag1*^−/−^ 2-3 weeks post naïve T cell transfer—were activated in round-bottom 96-well plates with mouse anti-CD3/anti-CD28 T cell expander beads (1 bead/cell; Life Technologies) in complete media containing 10 U/mL recombinant human (rh) IL-2 for 24 hr in the presence or absence of synthetic or endogenous CAR agonists (see ‘compound and tissue extracts’ below).

### Retroviral plasmids and transductions

shRNAmirs against mouse nuclear receptors were purchased (TransOMIC) or custom synthesized using the shERWOOD algorithm^41^. For cloning into an ametrine-expressing murine retroviral vector (LMPd) containing the enhanced miR-30 cassette^42,43^, shRNAmirs were PCR amplified using forward (5′-AGAAGGCTCGAGAAGGTATATTGC-3′) and reverse (5′-GCTCGAATTCTAGCCCCTTGAAGTC CGAGG-3′) primers containing XhoI and EcoR1 restriction sites, respectively. All retroviral constructs were confirmed by sequencing prior to use in cell culture experiments. Retroviral particles were produced by transfection of Platinum E (PLAT-E) cells with the TransIT-LT1 transfection reagent (Mirus) in Opti-MEM I reduced serum medium. Viral supernatants containing 10 μg/mL polybrene were used to transduce CD4^+^CD25^−^ T cells 24 hr post-activation (anti-CD3/anti-CD28; as above). Transductions were enhanced by centrifugation at 2000 rpm for 1 hr at room temperature, and incubation at 37 °C until 48 hr post-activation. Transduced cells were expanded in complete media containing 10 U/mL rhIL-2.

### Cell lines

PLAT-E cells, derived from the HEK-293 human embryonic kidney fibroblasts and engineered for improved retroviral packaging efficiency, were provided by M. Pipkin (Scripps Florida). All cell lines were tested to be mycoplasma free, and cultured in DMEM plus 10% FBS, 2 mM L-glutamine, 50 uM 2-mercaptoethanol, 1% HEPES, 0.1% gentamicin and 100u/ml Pen-Strep.

### T cell transfer colitis

For experiments using B6-derived wild-type or CAR-deficient (*Nr1i3*^−/−^) T cells, 0.5 × 10^6^ FACS-sorted naïve T cells (sorted as CD4^+^CD25^−^CD62L^hi^CD44^lo^ at Scripps Florida; CD4^+^CD45RB^hi^ at BCM) were injected intraperitoneally (i.p.) into syngeneic *Rag1*^−/−^ (at Scripps Florida) or *Rag2*^−/−^ (at BCM) recipients and analyzed between 2-6 weeks post-transfer. For mixed congenic T cell transfers, FACS-purified naïve T cells (CD4^+^CD25^−^CD62L^hi^CD44^lo^) from CD45.1 wild type and CD45.2 CAR-deficient (*Nr1i3*^−/−^), PXR-deficient (*Nr1i2*^−/−^), CAR- and PXR-deficient (*Nri12*^−/−^*Nr1i3*^−/−^) or *Il10*^−/−^ mice were mixed in a 1:1: ratio and transferred together (0.5 x 10^6^ total cells). For transfers of shRNAmir-expressing T cells, magnetically enriched CD4^+^CD25^−^ T cells from FVB/N (FVB) wild-type mice, activated and transduced as above, were expanded until day 5 in media containing rhIL-2 and transferred into syngeneic *Rag1*^−/−^ mice (0.5 x 10^6^ total cells). All *Rag1*^−/−^ recipients were weighed immediately prior to T cell transfer to determine baseline weight, and then weighed twice weekly after T cell transfer for the duration of the experiment. Mouse chow diets containing 2% Cholestyramine (CME) (Sigma-Aldrich) or 0.2% Cholic Acid (CA) (Sigma-Aldrich) and control diets were custom made (Teklad Envigo, Madison, WI) and fed to mice as follows: CME-supplemented diets were started 3 weeks after T cell transfer and continued for 3 weeks; cholic acid diet was started within 3 days post-T cell transfer and continued for 6 weeks (or until mice died). TCPOBOP (TC; Sigma-Aldrich) was initially reconstituted in sterile DMSO, stored at −20 °C, and diluted in sterile saline and sonicated immediately prior to injections. 3 mg/kg TC was injected intra-peritoneal (i.p.) weekly as indicated. Transferred *Rag1*^−/−^ or *Rag2*^−/−^ mice were euthanized upon losing 20% of pre-transfer baseline weight. All *Rag*^−/−^ mice receiving different donor T cell genotypes were co-housed to normalize microflora exposure.

### Anti-CD3-induced intestinal injury

Wild-type (B6) or CAR-deficient (B6.*Nr1i3*^−/−^) mice were injected i.p. with 15 ug of soluble, Ultra-LEAF purified anti-CD3 (clone: 145-2C11) or IgG isotype control (clone: HTK888) (BioLegend) twice over 48 hr. Animals were euthanized, and T cells analyzed 4 hr after the second injection.

### Histology

Colon (proximal, distal) or small intestine (proximal, mid, distal/ileum) sections (~ 1 cm) were cut from euthanized *Rag1*^−/−^ or *Rag2*^−/−^ mice 6 weeks post-T cell transfer. In some experiments, 10 cm segments of distal small intestine and whole colon were dissected from mice and fixed intact. All tissues were fixed in 10% neutral buffered formalin, embedded into paraffin blocks, cut for slides at 4-5 microns, and stained with hematoxylin and eosin (H&E). H&E-stained sections were analyzed and scored blindly by a pathologist with GI expertise using an Olympus BX41 microscope and imaged using an Olympus DP71 camera. Colons and ilea were histologically graded for inflammation severity using a combination of previously-reported grading models published by Kim, et al.^31^ and by Berg et al.^32^. The scheme published by Kim, et al grades 5 different descriptors which include crypt architecture (normal, 0 - severe crypt distortion with loss of entire crypts, 3), degree of inflammatory cell infiltration (normal, 0 – dense inflammatory infiltrate, 3), muscle thickening (base of crypt sits on the muscularis mucosae, 0 - marked muscle thickening present, 3), goblet cell depletion (absent, 0-present, 1) and crypt abscess (absent, 0-present, 1). The histological damage score is the sum of each individual score.

### Flow cytometry

Cell surface and intracellular FACS stains were performed at 4 °C for 30 minutes, washed with phosphate buffered saline (PBS) and acquired on a flow cytometer. Analysis of Rh123 efflux was performed as in^4^. Background Rh123 efflux was determined by the addition of the MDR1 antagonist, elacridar (10 nM), to Rh123-labelled cells prior to the 37 °C efflux step. Anti-mouse antibodies used for FACS analysis included: Alexa Fluor 700 anti-CD45, APC anti-CD45.1, BV711 anti-CD4, BV510 anti-CD25, BV650 anti-CD3, Percp-Cy5.5 anti-CD62L, PE-CY7 anti-CD44, BV605 anti-CD62L, PE anti-α4β7, Alexa Fluor700 anti-CD4, FITC anti-CD44, BV421 anti-CD44, e450 anti FOXP3, BV605 anti-TNF, Percp-Cy5.5 anti-Il-17a, BV711 anti-INFγ, PE anti-Il-4, PE-CY7 anti-IL-10, PE anti-Thy1.1, FITC anti-CD3, Percp-Cy5.5 anti-Thy1.1, PE anti-CD3, PE anti-TCRβ, APC anti-INFg, FITC anti-CD45.2, PE anti-α4β7 (from BioLegend); and BUV395 anti-CD3, PE-CF594 anti-CD25, FITC anti-Ki-67, PE-CF594 anti-RORγt, FITC anti-CD4, PE anti-CD45RB (from BD). Anti-human antibodies used for FACS analysis included: APC anti-CD3, PE anti-CD4, PE-Cy7 anti-CD45RO, BV711 anti-CD49a (integrin α4), APC-Fire 750 anti-integrin β7, BV421 anti-CCR9, and Percp-Cy5.5 anti-CCR7, BV605 anti-CCR2, PE anti-CRTH2, PE anti-CCR10, PE-Cy7 anti-CCR4, Percp-Cy5.5 anti-CXCR3, APC anti-CCR6, BV605 anti-CD4, PE-CF594 anti-CD25 (from BD). Vital dyes include: fixable viability eFluor^®^ 506, eFluor^®^ 660 and eFluor^®^ 780 (all from eBioscience). Rh123 and elacridar were purchased from Sigma-Aldrich. All FACS data was acquired on LSRII or FACSCanto II instruments (BD), and analyzed using FlowJo 9 or FlowJo 10 software (TreeStar, Inc.). (We’re probably missing a bunch).

### Cell sorting

Cells stained with cell-surface antibodies, as above, were passed through 70 μm nylon filters, resuspended in PBS plus 1% serum, and sorted on a FACS AriaII machine (BD Biosciences). Sorted cells were collected in serum-coated tubes containing PBS plus 50% serum. Gates used to sort MDR1+/− T cells, based on Rh123 efflux, were set using background Rh123 efflux in elacridar-treated cells. For human T cell sorts, Peripheral blood mononuclear cells (PBMC) were isolated using Ficoll-Plaque PLUS (GE Healthcare) from 25 mL of enriched buffy coats (OneBlood). CD4^+^ T cells were enriched using the Human total CD4 T cell Negative Isolation kit (EasySep), followed by enrichment of either effector/memory T cells (Human Memory CD4 T cell Enrichment kit; EasySep) or Treg cells (Human CD4^+^CD127^lo^CD49d^−^ Treg Enrichment Kit; EasySep) (all from StemCell Technologies).

Enriched cells were stained with anti-human FACS antibodies (listed above) for 20 minutes on ice. Stained cells were filtered through sterile 40 uM mesh filters and re-suspended in PBS with 5% FBS and 0.1% DNase. In cases where RNA was isolated after sorting, 100,000 cells were sorted into 200 uL PBS with 1 uM DTT and 5 uL RNase Inhibitor Cocktail (Takara); for *ex vivo* culture experiments, 0.4-1.2 ×10^6^ cells were sorted into complete T cell media.

### Pooled *in vivo* shRNAmir screen

Two independent pooled screens were performed. Briefly, PLAT-E cells were cultured in 96 well plates with 5 × 10^4^ per well in 100uL complete medium and transfected as described above. Magnetically enriched CD4^+^CD25^−^ T cells from spleens of 7-to 8-week old female FVB/N (FVB) mice were activated with anti-CD3 and anti-CD28 in 96 well plates and transduced 24 hr post-activation. Transduction efficiency of each individual shRNA was determined on day 4; transduced cells were pooled and FACS-sorted for ametrine^+^ on day 5 and adoptively transferred (i. p.) into 10 FVB.*Rag1*^−/−^ mice. An aliquot of sorted cells was saved for genomic DNA isolation and used for input reference. Six weeks post-transfer, live (viability dye^−^) transduced (ametrine^+^) Rh123^hi^ (MDR1^−^) or Rh123^lo^ (MDR1^+^) effector/memory T cells (Teff; CD4^+^CD25^−^CD62L^lo^CD44^hi^) were FACS-sorted from the spleen or small intestine lamina propria of FVB.*Rag1^−/−^* recipients. High quality genomic DNA was isolated using PureLink^®^ Genomic DNA Mini Kit (Invitrogen) and 100 ng of DNA was used for library preparation. gDNA derived from transduced and sorted T cells were quantified with Qubit DNA assay. 75ng of gDNA were used as template in duplicate reactions to add the Ion adapter sequences and barcodes. Based on previous data, 28 cycles of PCRs were used to amplify the libraries using primers with Ion P1 miR30 loop sequence (5’-CCTCTCTATGGGCAGTCGGTGAT<u>TACATCTGTGGCTTC-ACTA</u>-3’) and Ion A miR-30 (5’-CCATCTCATCCCTGCGTGTCTCCGACTCAGXXXXXXXXXX-<u>GCTCGAGAAGGTATATTGCT</u>-3’) sequences. The miR-30 loop (PI) and miR-30 (A) annealing sequences are underlined. The IonXpress 10 nt barcode is depicted with a string of X’s. Sequencing libraries were purified with 1.6X Ampure XP beads (Beckman Coulter), quantified with Qubit DNA HS assay (Invitrogen), and visualized on the Agilent 2100 Bioanalyzer (Agilent Technologies, Inc.). Individually-barcoded libraries were pooled at equimolar ratios and templated on to Ion spheres at 50 pM loading concentration using the Ion Chef (Life Technologies) with the Ion PI IC 200 kit. The templated Ion spheres (ISPs) were quantified using AlexaFluor sequence-specific probes provided in the Ion Spehere quality control kit (Life Technologies). The percent templated ISPs within 10-20% were taken forward to loading on the Ion PI V2 chips and then run on the Ion Proton with 200 bp reads. Libraries were sequenced using the Ion Torrent technology from Life Technologies following the manufacturer’s instructions. Sequencing reads were aligned to the reference library using BLAST with default settings and raw counts were normalized with DESeq2. Normalized reads of shRNAmirs displaying ≤ 10-fold change between input and *ex vivo* spleen samples were considered for downstream analysis. The relative enrichment or depletion of shRNAmirs was determined using median log_2_ fold-changes in shRNAmir abundances in MDR1^hi^ *vs*. MDR1^lo^ Teff cells. Median values for each gene target were calculated based on mean shRNAmir abundances determined in 2-indpeendent screens, each using cells recovered from pools of 10 spleens and siLP of transferred FVB.*Rag1*^−/−^ mice.

### Compounds and tissue extracts

10 or 20 uM 1,4-Bis-[2-(3,5-dichloropyridyloxy)]benzene, 3,3′,5,5′-Tetrachloro-1,4-bis(pyridyloxy) benzene (TC), 10 uM 5α-Androstan-3β-ol (And), 10 uM 5-Pregnen-3β-ol-20-one-16α-carbonitrile (PCN) (all from Sigma-Aldrich)—or serum, bile (from gallbladder), sterile soluble small intestine lumen content (siLC), or sterile soluble colon lumen content (cLC) from wild type (B6) mice—were added to mouse naïve or effector/memory (Teff) cells stimulated with anti-CD3/anti-CD28 antibodies as above. For preparation of mouse tissue extracts, mouse small intestinal lumen content (siLC) or colon lumen content (cLC) was extracted from whole tissue into a sterile tube. Contents were weighed, diluted with an equal volume of sterile PBS, vortexed vigorously for 30 sec, and then supernatants were collected after sequential centrifugation steps: (*i*) 10 min at 930 x *g*; and (*ii*) 10 min at 16 x *g*. Cleared supernatants were finally sterile-filtered using 0.22 μm filters and aliquots were frozen at −20° C. Serum was collected in EDTA coated tubes and centrifuged for 5 min at 2.4 x *g.* Due to small sample size, serum and gallbladder bile were used directly without filter sterilization after harvesting. Equal volumes of sterile vehicles (DMSO for TC, And; ethanol for PCN; PBS for sterile mouse content) served as negative controls. For human T cell culture experiments, healthy adult donor PBMC were FACS-sorted for the following subsets: (*i*) naïve CD4^+^ T cells (CD4^+^CD25^−^CD45RO^−^CCR7^hi^); (*ii*) Treg cells (CD4^+^CD25^hi^); (*iii*) α4^−^CCR9^−^ effector/memory cells (Teff; CD4^+^CD25^−^CD45RO^+^); and (*iv*) α4^+^CCR9^+^ effector/memory cells (Teff; CD4^+^CD25^−^ CD45RO^+^). Note that all α4^−^CCR9^−^ Teff cells are integrin β7^−^and all α4^+^CCR9^+^ Teff cells are integrin β7^+^. For all subsets, 30,000 purified cells were stimulated in round-bottom 96-well plates with human anti-CD3/anti-CD28 T cell expander beads (1 bead/cell; ThermoFisher) in complete media containing 10 U/mL rhIL-2 with or without 10 or 20 uM 6-(4-Chlorophenyl)imidazo[2,1-b][1,3]thiazole-5-carbaldehyde O-(3,4-dichlorobenzyl)oxime (CITCO) (Sigma-Aldrich); an equal volume of DMSO served as the negative control.

### qPCR

RNA was isolated from cultured or ex vivo-isolated cells using RNeasy Mini columns with on-column DNase treatment (Qiagen); RNA was used to synthesize cDNA via a high capacity cDNA reverse transcription kit (Life Technologies). Taqman qPCR was performed on a StepOnePlus real time PCR instrument (Life Technologies/Applied Biosystems) using commercial Taqman primer/probe sets (Life Technologies). Probes for mouse genes included: *Abcb1a* (Mm00607939_s1), *Nr1i3* (Mm01283981_g1), *Cyp2b10* (Mm01972453_s1), *Il10* (Mm01288386_m1) and *Actin b (Mm00607939_s1)*; probes for human genes included: *NR1I3* (Hs00901571_m1), *ABCB1* (Hs00184500_m1), *CYP2B6* (Hs04183483), *IL10* (Hs00961622_m1), and *ACTIN B* (Hs0160665_g1).

### Bioinformatics

#### ChIP-seq

Raw sequencing reads for CAR were downloaded from Gene Expression Omnibus (GSE112199)^17^, aligned to USC mm10 with Bowtie2^33^ and analyzed with MACS^34^ using base settings. Biological replicate reads files were merged into a single file and bigwig files were generated and visualized with Integrated Genome Viewer (IGV)^35^. Peaks were filtered to remove reads with alternative annotations, mitochondrial DNA, or blacklist regions in R using GenomeInfoDb and GenomicRanges package.

#### RNA-seq

Next-generation RNA-sequencing (RNA-seq) was performed on FACS-sorted B6 wild type and CAR-deficient effector/memory T cells (Teff cells: viability dye^−^CD45^+^CD3^+^CD4^+^CD25^−^CD44^hi^) from spleen, small intestinal lamina propria, and colon lamina propria of *Rag1*^−/−^ mice injected 3-weeks prior with congenic mixtures of CD45.1 wild type and CD45.2 *Nr1i3*^−/−^ naïve T cells, approximately 500 sorted cells were processed directly to generate cDNA using the Clontech SMART-Seq v4 Ultra Low Input Kit (Clontech, Inc.) on three biologically-independent replicates. The generated cDNA was size selected using beads to enrich for fragments > 400 bp. The enriched cDNA was converted to Illumina-compatible libraries using the NEBNext Ultra II DNA kit (New England Biolabs, Inc.) using 1ng input. Final libraries were validated on the Agilent 2100 bioanalyzer DNA chips and quantified on the Qubit 2.0 fluorometer (Invitrogen, Life Technologies). Barcoded libraries were pooled at equimolar ratios and sequenced using single-end 75 bp reads on a NextSeq 500 instrument (Illumina). Raw sequencing reads (fastq files) were mapped to the mm10 transcriptome and transcript abundance in terms of Transcripts Per Million (TPM) were quantified using Salmon^48^. PCA was performed and projected in R-studio. Differentially expressed genes (DEG) were determined using DESeq2 (*P* < .05) for CAR-deficient (B6.*Nr1i3*^−/−^) *vs.* wild type (B6) Teff cells from spleen (296 up; 285 down), siLP (472 up; 523 down), or cLP (350 up; 228 down) and log_2_ fold-change was used as the ranking metric to generate input ranked lists for gene set enrichment analysis (GSEA) (https://www.gsea-msigdb.org/gsea/index.jsp); these genes were compared against both customized and curated gene sets (the latter from the Molecular Signature Database (MSigDB)) for enrichment—quantified as normalized enrichment score (NES)—and visualized using ggplot2 package in R. For GSEA summary plots (Fig 2d, 3b; Extended Data Fig. 3b) circle sizes indicate significance (−log_10_ Padj values). Red/blue coloring indicates enrichment within genes up/down, respectively, in CAR-deficient *vs*. wild type cells, based on normalized enrichment score (NES). Differentially expressed genes of wild type (B6) Teff cells from the spleen, siLP, or cLP determined by DESeq2 were used to generate tissue-specific Teff gene sets: (*i*) up in B6 spleen Teff, genes selectively expressed in spleen *vs*. either siLP or cLP wild type (B6) Teff cells; (*ii*) up in B6 siLP Teff, genes selectively expressed in siLP *vs*. either spleen or cLP wild type (B6) Teff cells; and (*iii*) up in B6 cLP Teff, genes selectively expressed in cLP *vs*. either spleen or siLP wild type (B6) Teff cells. RNA-seq data of pharmacological activation of CAR or PXR in hepatocytes *in vivo* from mice treated with the CAR agonist, TCPOBOP (TC), the PXR agonist, PCN, or vehicle (corn oil) (GSE104734)^16^ were analyzed to generate the gene sets: Up in Hep + TC, genes selectively induced by the CAR agonist, TCPOBOP (TC), compared with either vehicle (corn oil) or the PXR agonist, PCN, in hepatocytes from mice treated with compounds *in vivo*; and Up in Hep + PCN, genes selectively induced by the PXR agonist, PCN, compared with either vehicle (corn oil) or the CAR agonist, TC, in hepatocytes from mice treated with compounds *in vivo*. Differential gene expression of *in vitro*-differentiated Tr1 (GSE92940)^22^ and Th17 cells (GSE21670)^36^ were determined using the Limma package in R (for microarray data)^37^ to generate the gene sets: Tr1-signature, genes selectively expressed in *in vitro*-differentiated Tr1 cells, compared with non-polarizing conditions; and Th17-signature, genes selectively expressed in *in vitro*-differentiated Th17 cells, compared with non-polarizing conditions. Th1-signature, Th2-signature, induced (i)Treg-signature (GSE14308)^38^, or T follicular helper (Tfh)-signature (GSE21379)^39^, genes selectively induced in these *vs*. other T cell subsets, as curated on MSigDB (https://www.gsea-msigdb.org/gsea/msigdb/index.jsp).

### TR-FRET co-regulator recruitment assay

The DNA sequences encoding mouse (m)CAR ligand-binding domain (LBD; residues 109 – 358) were amplified by PCR reaction and inserted into modified pET24b vectors to produce pET24b-mCAR-LBD. pACYC-Duet1-RXR-LBD, an expression plasmid for untagged human (h)RXRα LBD was provided by Dr. Eric Xu^40^. Purification of the mCAR-hRXRα LBD heterodimer, as well as hRXRα homodimer, was achieved by nickel-affinity chromatography, followed by size-exclusion chromatography in an Akta explorer FPLC (GE Healthcare). Briefly, pET24b-mCAR-LBD and pACYC-Duet1-RXR-LBD were co-transformed into BL21 (DE3) for mCAR-hRXRα heterodimer and pET46-RXRα-LBD was transformed into BL21 (DE3) for RXRα homodimer. The cells were grown in 4 x 900 mL of LB media at 37 °C until the OD600 reached a value of 0.6–0.7. Overexpression was induced by 0.3 mM of IPTG and the cells were grown further for 22 hr at 18 °C. The harvested cells were resuspended in sonication buffer (500 mM NaCl, 10 mM HEPES, 10 mM imidazole, pH 8.0, and 10% glycerol), sonicated on an ice-water bath for 20 min at 18 W output, and centrifuged for 25 min at 50,000 x g. The proteins were isolated from the sonicated supernatant by applying to a 2 mL His Select column and eluted with linear gradient from 10 mM to 300 mM imidazole in sonication buffer. The elution fractions containing the proteins concentrated while exchanging buffer to gel filtration buffer (300 mM NaCl, 20 mM HEPES, 1 mM DTT, 5 % glycerol). The proteins were purified further by gel filtrations through a Superdex 200 26/60 column (GE Healthcare) equilibrated with gel filtration buffer. Fractions containing the proteins were pooled and concentrated to ~ 8 mg/mL each with 30 kDa cutoff ultrafiltration units (Millipore). Time-resolved fluorescence resonance energy transfer (TR-FRET) assays were performed in low-volume black 384-well plates (Greiner) using 23◻μL final well volume. Each well contained the following components in TR-FRET buffer (20 mM KH_2_PO_4_/K_2_HPO_4_, pH 8, 50 mM KCl, 5 mM TCEP, 0.005% Tween 20): 4 nM 6xHis-CAR/RXRα LBD heterodimer or 6xHis-RXRα/RXRα homodimer LBD, 1◻nM LanthaScreen Elite Tb-anti-His Antibody (ThermoFisher #PV5895), and 400 nM FITC-labeled PGC1α peptide (residues 137–155, EAEEPSLLKKLLLAPANTQ, containing an N-terminal FITC label with a six-carbon linker, synthesized by Lifetein). Pure ligand (TC, 9-*cis* RA) or tissue extracts (see above) were prepared via serial dilution in vehicle (DMSO or PBS, respectively), and added to the wells along with vehicle control. Plates were incubated at 25 °C for 1 hr and fluorescence was measured using a BioTek Synergy Neo plate reader (Promega). The terbium (Tb) donor was excited at 340 nm, its emission was monitored at 495 nm, and emission of the FITC acceptor was monitored at 520 nm. Data were plotted as 520/340 nM rations using Prism software (GraphPad); TC data were fit to a sigmoidal dose response curve equation.

### Statistical Analyses

Statistical analyses were performed using Prism (GraphPad). *P* values were determined by paired or unpaired student’s *t* tests, Log-rank test, one-way ANOVA, and two-way ANOVA analyses as appropriate and as listed throughout the Figure legends. Statistical significance of differences (* P < 0.05, **P < 0.01, ***P < 0.001, ****P < 0.0001) are specified throughout the Figure legends. Unless otherwise noted in legends, data are shown as mean values ± S.E.M.

### Reporting summary

Further details regarding research design is available in the Nature Research Reporting Summary linked to this paper.

## Data availability

RNA-seq data for wild type and CAR-deficient effector CD4^+^ T cells from spleen, small intestine lamina propria or colon lamina propria of congenically co-transferred *Rag1*^−/−^ mice, as well as from human peripheral blood α4^+^β7^+^CCR9^+^ memory CD4^+^ T cells stimulated *ex vivo* in the presence or absence of the human CAR agonist, CITCO, are available on the NCBI Gene Expression Omnibus (GEO) repository (accession ID: GSE149220).

## Code availability

No proprietary code was written or used for data analyses in this study.

## Acknowledgements

The authors thank Core Facility staff at Scripps Florida and Baylor College of Medicine for technical support, and Drs. Paul Dawson and Anjana Rao for critical discussions. This work was supported by Scripps Florida via the State of Florida (M.S.S.), the R.P. Doherty Jr.–Welch Chair in Science Q◻0022 at Baylor College of Medicine (D.D.M.), National Institute of Health grants R21AI119728 (M.S.S.), R01AI118931-01 (M.S.S.), U19AI109976 (M.E.P), P01AI145815 (M.E.P.), and a Senior Research Award from the Crohn’s and Colitis Foundation (M.S.S.).

## Author contributions

Study design: M-L.C., X.H., H.W., J.S., H.P., B.F., L.A.S., D.J.K., A.R-P., D.A.S., C.T.W., M.E.P., D.D.M., and M.S.S. Data generation: M-L.C., X.H., H.W., C.H., Y.L., J.S., A.E., and S.A.M. Bioinformatics: M-L.C., G.W., A.R-P., and M.E.P. Manuscript: M-L.C., X.H., C.H., A.E., D.J.K., A. R-P., C.T.W., M.E.P., D.D.M., and M.S.S. Principal Investigators: L.A.S., D.J.K., A.R-P., D.A.S., C.T.W., M.E.P., D.D.M., and M.S.S.

## Competing interests

M.S.S. is a consultant to Sigilon Therapeutics and Sage Therapeutics.

## Additional information

**Supplementary information** is available for this paper.

## Extended Data Figure Legends

**Extended Data Figure 1. Nuclear receptor-dependent regulation of effector T cell persistence and MDR1 expression *in vivo*. (a)** *Top*, abundance of shRNAmirs in *ex vivo*-isolated spleen and *in vitro*-transduced (input) Teff cells. shRNAmirs with ≤ 1 normalized read in both *ex vivo* spleen and input Teff cell pools were considered ‘poorly represented’ (highlighted green). Well-represented shRNAmirs displaying ≤ 10-fold change between *ex vivo* spleen and input Teff cell pools (between blue lines) were considered for downstream analysis. *Bottom*, abundance of shRNAmirs, filtered for minimal effects on *in vivo* Teff cell persistence, in *ex vivo*-isolated MDR1^hi^ (Rh123^lo^) and MDR1^lo^ (Rh123^hi^) siLP Teff cells. **(b)** Log_2_ fold-change in abundance (± SEM) of shRNAmirs against *Cd19* (*n* = 3), *Abcb1a* (*n* = 2), *Nr1i3* (*n* = 5), *Thra* (*n* = 6), and *Esrra* (*n* = 3) in FVB wild type Rh123^lo^ (MDR1^hi^) *vs*. Rh123^hi^ (MDR1^lo^) effector/memory T cells (Teff; sorted as in Fig. 1a) recovered from spleens or small intestine lamina propria (siLP) of transferred FVB.*Rag1*^−/−^ mice. (a-b) Data incorporates shRNAmir abundance, determined by DNA-seq, in 2-indpeendent screens using pooled spleens and siLP from 10 transferred FVB.*Rag1*^−/−^ mice per screen. **(c)** *Ex vivo* Rh123 efflux, determined by flow cytometry, in FVB wild type Teff cells expressing a control shRNAmir against CD8 (*shCD8a*) or 1 of 5-independent shRNAmirs against CAR (*shNr1i3*) isolated from spleens of transferred FVB.*Rag1*^−/−^ mice 6-weeks post-transfer. Rh123 efflux in transduced (Ametrine pos.; blue) cells is overlaid with that in congenically-transferred untransduced (Ametrine neg.; red) Teff cells from the same mouse. Background Rh123 efflux in untransduced Teff cells treated with the MDR1 inhibitor, elacridar, is shown in gray. Representative of 63 mice analyzed over 3-independent experiments. **(d)** Mean normalized *ex vivo* Rh123 efflux (± SEM) in FVB wild type spleen Teff cells expressing control (*shCd8a*; *n* = 11) or CAR-targeting (shNr1i3) shRNAmirs; *shNr1i3.1* (*n* = 10), *shNr1i3.2* (*n* = 10), *shNr1i3.3* (*n* = 12), *shNr1i3.4* (*n* = 10), *shNr1i3.5* (*n* = 10), determined by flow cytometry as in (c). Rh123 efflux was normalized to control *shCd8a*-expressing Teff cells after calculating the change (Δ) in Rh123 mean fluorescence intensity (MFI) between congenically-transferred transduced (ametrine pos.) *vs*. untransduced (ametrine neg.) Teff cells. * *P* < .05, **** *P* < .0001, One-way ANOVA with Dunnett’s correction for multiple comparisons. **(e)** Mean relative *Abcb1a*, *Nr1i3*, and *Cyp2b10* expression (± SEM), determined by qPCR, in FVB spleen Teff cells FACS-sorted from FVB.*Rag1*^−/−^ recipient mice expressing either a negative control shRNAmir against CD8 (*shCd8a*; *n* = 8), or the indicated shRNAmirs against CAR (*shNr1i3*s); *shNr1i3.1* (*n* = 8), *shNr1i3.2* (*n* = 8), *shNr1i3.3* (*n* = 8), *shNr1i3.4* (*n* = 8), *shNr1i3.5* (*n* = 8).* *P* < .05, ** *P* < .01, *** *P* < .001, **** *P* < .0001, One-way ANOVA with Tukey’s correction for multiple comparisons. **(f)** Median log_2_ fold change in shRNAmir abundance between FVB wild type *ex vivo*-isolated spleen *vs*. *in vitro*-transduced (input) Teff cells. (a, d) shRNAmir abundance reflects the mean number of normalized reads, by DNA-seq, in the indicated Teff subsets obtained in 2-independent screens, each using cells transferred into 10 FVB.*Rag1*^−/−^ mice. **(g)** *Ex vivo* Rh123 efflux, determined by flow cytometry, in CD45.1 wild type (B6; red) or CD45.2 CAR-deficient (B6.*Nr1i3*^−/−^), PXR-deficient (B6.*Nr1i2*^−/−^) or CAR/PXR double-deficient (B6.*Nr1i3*^−/−^*Nr1i2*^−/−^) effector/memory T cells (Teff; gated as in Extended Data Fig. 6a; blue) isolated from spleens of B6.*Rag1*^−/−^ mice 6-weeks post-naïve T cell congenic co-transfer. Background Rh123 efflux in CD45.1 B6 Teff cells treated with the MDR1 inhibitor, elacridar, is shown in gray. Representative of a total of 22 mice analyzed over two-independent T cell transfer experiments. **(h)** Mean normalized Rh123 efflux (± SEM) in congenically-transferred CD45.1 wild type (B6; *n* = 7) or CD45.2 CAR-deficient (B6.*Nr1i3*^−/−^; *n* = 7), PXR-deficient (B6.*Nr1i2*^−/−^; *n* = 7) or CAR/PXR double-deficient (B6.*Nr1i3*^−/−^*Nr1i2*^−/−^; *n* = 7) spleen Teff cells, determined by flow cytometry as in (g). * *P* < .05, One-way ANOVA with Tukey’s correction for multiple comparisons. **(i)** Mean relative *Abcb1a* expression (± SEM), determined by *ex vivo* qPCR, in CD45.1 wild type (B6; *n* = 5) or CD45.2 CAR-deficient (B6.*Nr1i3*^−/−^; *n* = 5), PXR-deficient (B6.*Nr1i2*^−/−^; *n* = 4) or CAR/PXR double-deficient (B6.*Nr1i3*^−/−^*Nr1i2*^−/−^; *n* = 5) spleen Teff cells (sorted as in Extended Data Fig. 6a) from spleens of congenically-transferred B6.*Rag1*^−/−^ as in (a). * *P* < .05, One-way ANOVA with Tukey’s correction for multiple comparisons.

**Extended Data Figure 2. Inhibition of bile acid reabsorption rescues ileitis induced by CAR-deficient T cells in reconstituted *Rag*^−/−^ mice. (a)** Mean weight loss (+SEM) of co-housed B6.*Rag2*^−/−^ mice transplanted with wild type (B6; blue; *n* = 15) or CAR-deficient (B6.*Nr1i3*^−/−^; red; *n* = 13) naïve CD4^+^ T cells and treated with 2% (w:w) cholestyramine (CME) beginning at 3-weeks post-T cell transfer. NS, not significant. **(b)** *Top*, H&E-stained sections of colons or terminal ilea from B6.*Rag2*^−/−^ mice reconstituted with wild type or CAR-deficient T cells and treated +/− CME as in (a). Representative of 12 mice/group. *Bottom*, mean histology scores (± SEM; *n* = 12) for colons or terminal ilea as in (a). NS, not significant. **(c)** Mean weight loss (+SEM) of co-housed B6.*Rag1*^−/−^ mice with or without the Apical sodium-dependent bile acid transporter (Asbt; gene symbol *Slc10a2*) after transplantation with wild type (B6; blue) or CAR-deficient (B6.*Nr1i3*^−/−^; red) naïve CD4^+^ T cells. **(d)** *Top*, H&E-stained sections of terminal ilea or colons from control or Asbt-deficient B6.*Rag1*^−/−^ mice reconstituted with wild type or CAR-deficient T cells as in (c). Representative of 5 mice/group. *Bottom*, mean histology scores (± SEM; *n* = 5) for colons or terminal ilea as above. * *P* < .05, ** *P* < .01, *** *P* < .001, One-way ANOVA with Tukey’s correction for multiple comparisons. NS, not significant.

**Extended Data Figure 3. Shared features of CAR-dependent gene expression in mucosal T cells and hepatocytes. (a)** Overlap, presented as Venn diagrams, between genes induced in B6 wild type mouse hepatocytes by *in vivo* treatment with either the mouse CAR agonist, TCPOBOP (TC) or the mouse PXR agonist, PCN, relative to vehicle (CO, corn oil). **(b)** Enrichment of genes induced by TC, but not PCN, treatment in mouse hepatocytes (as in [a]), within those reduced in CAR-deficient (B6.*Nr1i3*^−/−^) *vs*. wild type (B6) siLP Teff cells from week-3 congenically co-transferred *Rag1*^−/−^ mice (as in Fig. 2a-c). **(c)** Differential gene expression, determined by DEseq2 and shown as a volcano plot, between CAR-deficient (B6.*Nr1i3*^−/−^) and wild type (B6) siLP Teff cells re-isolated from transferred B6.*Rag1*^−/−^ mice, as in Fig. 2a. Genes induced by TC, but not PCN, treatment in mouse hepatocytes (as in [a]; purple), bound by CAR in ChIP-seq analysis of hepatocytes from TC-treated mice (blue), or both (red) are highlighted. Chi-square *P* values are indicated. **(d)** CAR-occupancy, determined by ChIP-seq, at representative loci whose expression is regulated by CAR in both mucosal T cells and hepatocytes within mouse hepatocytes ectopically expressing epitope-tagged mouse (m) or human (h) CAR proteins and re-isolated from mice after treatment with the mCAR agonist, TCPOBOP (TC), or the hCAR agonist, CITCO. * *P* < 0.00001; significant binding peaks were called in MACS using base settings.

**Extended Data Figure 4. CAR promotes effector T cell persistence in the presence of small intestinal bile acids. (a)** Percentages of live CD44^hi^ wild type (B6; CD45.1^+^; blue) or CAR-deficient (B6.*Nr1i3*^−/−^; CD45.1^−^; red) effector/memory (Teff) cells, determined by flow cytometry and gated as in Extended Data Fig. 6a, in tissues of reconstituted B6.*Rag1*^−/−^ mice over time. Numbers indicate percentages; representative of 5 mice per tissue and timepoint. **(b)** Fitness, defined as mean log_2_ fold-change (F.C.) of CAR-deficient (B6.*Nr1i3*^−/−^) *vs*. wild type (B6) Teff cell percentages (± SEM; *n* = 5) in tissues of congenically co-transferred *Rag1*^−/−^ mice over time, determined by flow cytometry as in (a). **(c)** Percentage of wild type (B6, CD45.1^+^; blue) and CAR-deficient (B6.*Nr1i3*^−/−^, CD45.1^−^; red) naïve (CD62L^hi^) CD4^+^ T cells after sorting and mixing, and prior to *in vivo* transfer into *Rag1*^−/−^ mice (input Tnaive); representative of 3 mixtures used for analyzing resulting Teff cells at 2- 4- or 6-weeks post-transfer. **(d)** Equal numbers of CD45.1 wild type (B6; blue) and CD45.2 CAR-deficient (B6.*Nr1i3*^−/−^; red) naïve CD4^+^ T cells were transferred together into co-housed *Rag1*^−/−^ mice with or without the ileal bile acid reuptake transporter, Asbt (gene symbol *Slc10a2*). Resulting effector (Teff) cells from small intestine lamina propria (siLP) were analyzed 2-weeks post-T cell transfer via flow cytometry. **(e)** Percentages of live CD44^hi^ wild type (B6; CD45.1^+^; blue) or CAR-deficient (B6.*Nr1i3*^−/−^; CD45.1^−^; red) effector/memory (Teff) cells, determined by flow cytometry and gated as in Extended Data Fig. 6a, in siLP of week-2 reconstituted B6.*Rag1*^−/−^ mice. Numbers indicate percentages; representative of 8-10 mice analyzed over two-independent experiments. **(f)** Mean absolute numbers (± SEM) of live CD45.1 wild type (B6; *left*) or CD45.2 CAR-deficient (B6.*Nr1i3*^−/−^; *right*) Teff cells, determined by *ex vivo* flow cytometry as in (e), from siLP 2-weeks after mixed T cell transfer into control (Asbt^+/+^; blue; *n* = 8) or Asbt-deficient (Asbt^−/−^; red; *n* = 10) *Rag1*^−/−^ recipients. Fold-changes in cell numbers recovered from Asbt-deficient *vs*. control recipients, as well as *P* values (two-tailed unpaired student’s t test) are indicated.

**Extended Data Figure 5. Preferential CAR expression and function in human effector/memory T cells expressing small bowel homing receptors. (a)** FACS-based identification of human CD4^+^ T cell subsets in PBMC from healthy adult human donors. Expression of integrin α4 (α4 int.) in gated naïve (gray), T regulatory (Treg; blue), or effector/memory (Teff; red) T cells is shown at right. **(b)** Expression of integrin β7 (β7 int.) and CCR9 in total naïve CD4^+^ T cells, or in α4 int.+/− Treg or Teff subsets (gated as in (a)). Representative of 13-independent experiments using PBMC from different donors. **(c)** Percentages (%) of α4^+^β7^+^CCR9^+^ Tnaive, Treg, or Teff cells, determined by flow cytometry as in (a-b). Individual data points for the 13 independent experiments are shown and connected by grey lines. ** *P* < .01, One-way ANOVA with Holm-Sidak’s correction for multiple comparisons. **(d)** *Ex vivo* Rh123 efflux in CD4^+^ T cell subsets (gated as in a-b) in the presence (gray) or absence (red) of the selective MDR1 inhibitor, elacridar. Representative of 8 experiments. **(e)** Mean percentages (± SEM; *n* = 7) of Rh123^lo^ (MDR1^+^) Teff subsets, determined by flow cytometry as in (d). * *P* < .05, ** *P* < .01, *** *P* < .001, One-way ANOVA with Tukey’s correction for multiple comparisons. **(f)** Mean (± SEM) *ex vivo* expression, determined by qPCR, of CAR/*NR1I3* (*n* = 12), MDR1/*ABCB1* (*n* = 12) or *CYP2B6* (*n* = 10) in α4^−^β7^−^CCR9^−^ or α4^+^β7^+^CCR9^+^ Tnaive, Treg or Teff cells, FACS-sorted as in (a-b). (e-f) * *P* < .05, ** *P* < .01, One-way ANOVA with Tukey’s correction for multiple comparisons. **(g)** Mean relative *CYP2B6* expression (± SEM; *n* = 5), determined by qPCR, in CD4^+^ T cell subsets (as in (f)) activated *ex vivo* with anti-CD3/anti-CD28 antibodies in the presence or absence of titrating concentrations of the human CAR agonist, CITCO. Gene expression was analyzed 24 hr post-activation. *** *P* < .001, Two-way ANOVA. **(h)** Mean normalized MDR1/*ABCB1* or *CYP2B6* expression (± SEM), determined by RNA-seq and presented as transcripts per million (TPM), in FACS-sorted α4^+^β7^+^CCR9^+^ Teff cells stimulated *in vitro* (anti-CD3/anti-CD28) for 24 hr in the presence or absence of CITCO. Data from 4 replicate RNA-seq experiments are shown; ** *P* < .001, paired two-tailed student’s *t* test. **(i)** Identification of CD4^+^ naive (Tnaive; CD25^−^CD45RO^−^; grey) or effector/memory (Teff; CD25^−^CD45RO^+^; red) cells, by flow cytometry, from healthy adult human PBMC. For improved purity of Th1, Th2, Th17 and Th17.1 cells, CCR10-expressing Th22 cells were excluded. CCR6 expression in Tnaive (grey) or non-Th22 Teff cells (red) is shown at right; CCR6^+^ or CCR6^−^ Teff cells were gated to enrich for Th17 or non-Th17 lineages, respectively. **(j)** Expression of CCR4 and CXCR3 in CCR6^−^ (non-Th17; *left*) or CCR6^+^ (Th17; *right*) Teff cells identifies enriched CCR6^−^CCR4^lo^CXCR3^hi^ (Th1; orange), CCR6^−^CCR4^hi^CXCR3^lo^ (Th2; blue), CCR6^+^CCR4^hi^CXCR3^lo^ (Th17; green), and CCR6^+^CCR4^lo^CXCR3^hi^ (Th17.1; red) subsets. **(k)** Expression of integrin α4 (α4 int.; *top*) in Th2, Th1, Th17 and Th17.1 human Teff cells gated as in (a-b). Expression of integrin β7 (β7 int.) and CCR9 within α4 int^−^ (*middle*) or α4 int^+^ (*bottom*) Th2, Th1, Th17 or Th17.1 cells gated as above. (a-c) Representative of 9-independent experiments using PBMC from different healthy adult donors. **(l)** Percentages (*n* = 9) of α4^+^β7^+^CCR9^+^ cells within *ex vivo* Th1, Th2, Th17, or Th17.1 Teff cells gated as in (a-c). Data from independent donors are connected by black lines. **(m)** MDR1-dependent Rh123 efflux in the indicated Th1, Th2, Th17, or Th17.1 Teff subsets gated based on expression of α4 int., β7 int., and/or CCR9 in the presence (grey) or absence (red) of elacridar. Representative of 8 independent experiments using PBMC from different donors. **(n)** Mean percentages (± SEM; *n* = 8) of Rh123^lo^ (MDR1^+^) cells within Th1, Th2, Th17, or Th17.1 Teff subsets gated based on expression of α4 int., β7 int., and/or CCR9 as in (e). * *P* < .05, ** *P* < .01, *** *P* < .001, One-way ANOVA with Tukey’s correction for multiple comparisons. ND, not detectable; NS, not significant.

**Extended Data Figure 6. TCPOBOP promotes CAR-dependent gene expression in *ex vivo*-isolated effector T cells. (a)** *Top left,* equal numbers of CD45.1 wild type (B6; blue) and CD45.2 CAR-deficient (B6.*Nr1i3*^−/−^; red) naïve CD4^+^ T cells were transferred together into B6.*Rag1*^−/−^ mice. Resulting effector (Teff) cells were FACS-purified from spleen after 3 weeks. *Right*, sequential gating strategy for re-isolating wild type and CD45.2 CAR-deficient spleen Teff cells is shown. *Bottom left*, mean relative *Abcb1a*, *Cyp2b10*, or *Il10* expression (± SEM; *n* = 4), determined by qPCR, in *ex vivo*-isolated wild type (B6) or CAR-deficient (B6.*Nr1i3*^−/−^) spleen Teff cells. These cells were used for *ex vivo* cell culture experiments in the presence or absence of small molecule ligands ([b-c] below). * *P* < .05, ** *P* < .01, paired two-tailed student’s *t* test. **(b)** Mean relative expression (± SEM) of *Abcb1a* (*n* = 4), *Cyp2b10* (*n* = 4), or *Il10* (*n* = 3), determined by qPCR, in wild type (B6) or CAR-deficient (B6.*Nr1i3*^−/−^) Teff cells isolated from transferred *Rag1*^−/−^ mice (as in [a]), and stimulated *ex vivo* with anti-CD3/anti-CD28 antibodies (for 24 hr) in the presence or absence of the mouse (m)CAR agonist, TCPOBOP (TC; 10 μM), the mCAR inverse agonist, 5α-Androstan-3β-ol (And; 10 μM), or both. ** *P* < .01, *** *P* < .001, **** *P* < .0001, one-way ANOVA with Tukey’s correction for multiple comparisons. **(c)** Mean relative *Abcb1a*, *Cyp2b10*, or *Il10* expression (± SEM; *n* = 5), determined by qPCR, in wild type (B6) or CAR-deficient (B6.*Nr1i3*^−/−^) Teff cells isolated and stimulated as in (a-b) in the presence or absence of TC (10 μM) or the mouse PXR agonist, PCN (10 μM). Data are presented as fold-change in mRNA abundance relative to vehicle-treated cells (DMSO for TC; ethanol for PCN). **** *P* < .0001, one-way ANOVA with Dunnett’s correction for multiple comparisons. NS, not significant.

**Extended Data Figure 7. Characteristics of endogenous intestinal metabolites that activate the CAR ligand-binding domain. (a)** Mean activation (± SEM; triplicate samples) of recombinant human (h)RXRα ligand-binding domain (LBD) homodimers, determined by time-resolved fluorescence resonance energy transfer (TR-FRET), in the presence of the mCAR agonist, TCPOBOP (TC; blue) or the hRXRα agonist, 9-*cis* retinoic acid (RA; red). Median effective concentration (EC_50_) of 9-*cis* RA-dependent hRXRα LBD homodimer activation is indicated. Representative of more than 5-independent experiments. **(b)** Mean activation (± SEM; *n* = 3) of hRXRα LBD homodimers, determined by TR-FRET as in (a), in the presence of titrating concentrations of siLC, bile, cLC or serum from wild type B6 mice. * *P* < .05, **** *P* < .0001, one-way ANOVA with Tukey’s correction for multiple comparisons. NS, not significant. **(c)** Mean activation (± SEM; *n* = 3) of CAR:RXR LBD heterodimers, determined by TR-FRET, in the presence of titrating concentrations of siLC isolated from conventionally-housed (Conv) or germ-free (GF) wild type B6 mice pre-treated with or without cholestyramine (CME) to deplete free bile acids. *** *P* < .001, **** *P* < .0001, One-way ANOVA with Dunnett’s correction for multiple comparisons. (a-c) The bars for each tissue extract indicate dilution series *(left to right)*: (1) diluent (PBS) alone; (2) 0.01%, (3) 0.1%, and (4) 1%. Data are shown from 3-independent experiments using extracts from different wild type mice, with each concentration from each individual mouse run in triplicate. **(d)** Mean TR-FRET signals (± SEM; *n* = 3) of CAR:RXR LBD heterodimers in the presence of titrating concentrations of individual bile acid (BA) species. NS, not significant, one-way ANOVA with Dunnett’s correction for multiple comparisons. The bars for BAs indicate concentrations *(left to right)*: (1) vehicle (DMSO); (2) 10 μM; (3) 100 μM; and (4) 1000 μM. Data are shown from 3-independent experiments, where each BA concentration was run in triplicate.

**Extended Data Figure 8. CAR promotes IL-10 expression in mucosal Teff cells and regulates Tr1 and Th17 cell development in the small intestine. (a)** Equal numbers of CD45.1 wild type (B6; blue) and CD45.2 CAR-deficient (B6.*Nr1i3*^−/−^; red) naïve CD4^+^ T cells were transferred together into *Rag1*^−/−^ mice. Resulting effector (Teff) cells were analyzed—using surface and intracellular flow cytometry after *ex vivo*-stimulation with phorbol myristate acetate (PMA) and ionomycin—at 2-4- and 6-weeks from spleen, mesenteric lymph node (MLN), small intestine lamina propria (siLP), or colon lamina propria (cLP). Gating hierarchy is shown from a representative sample of MLN mononuclear cells at 2-weeks post-T cell transfer. **(b)** Intracellular IL-10 and IFNγ expression, determined by flow cytometry, in wild type (B6, blue; *left*) or CAR-deficient (B6.*Nr1i3*^−/−^, red; *right*) non-Th17 Teff cells, gated as in (a), from tissues of T cell-reconstituted B6.*Rag1*^−/−^ mice over time. Numbers indicate percentages; representative of 5 mice per tissue and time point. Mean percentages **(c)** or numbers **(d)** (± SEM; *n* = 5) of IL-10-expressing wild type (B6, *left*) or CAR-deficient (B6.*Nr1i3*^−/−^, *right*) Teff cells, determined by *ex vivo* flow cytometry as in (a-b), from tissues of transferred B6.*Rag1*^−/−^ mice over time. **(e)** Specificity of IL-10 intracellular staining, as validated by analysis of IL-10 production by CD45.1 wild type (B6; blue) or CD45.2 *Il10*^−/−^ (red) Teff cells isolated from spleen or siLP of congenically co-transferred *Rag1*^−/−^ mice. Representative of 6 mice analyzed over 2-independent experiments. **(f)** Expression of RORγt and IL-17A, determined by intracellular FACS analysis as in Extended Data Figure 8a, in wild type (B6) or CAR-deficient (B6.*Nr1i3*^−/−^) CD4^+^ effector/memory (Teff) cells from tissues of reconstituted *Rag1*^−/−^ mice 2-weeks post-mixed T cell transfer. Numbers indicate percentages; representative of 5 mice per tissue and time point. **(g)** Mean percentages of (± SEM; *n* = 5) wild type (B6; blue) or CAR-deficient (B6.*Nr1i3*^−/−^; red) RORγt^+^IL-17A^−^ Teff cells, determined by intracellular flow cytometry as in (a). * *P* < .05, paired two-tailed student’s *t* test. **(h)** Expression of RORγt and IL-17A, determined by intracellular FACS analysis, in wild type (B6) or *Il10* ^−/−^ Teff cells from tissues of reconstituted *Rag1*^−/−^ mice 2-weeks post-mixed T cell transfer. Numbers indicate percentages; representative of 5 mice per tissue and time point. **(i)** Mean percentages of (± SEM; *n* = 7) wild type (B6; blue) or *Il10* ^−/−^ (red) RORγt^+^IL-17A^−^ Teff cells, determined by intracellular flow cytometry as in (c). * *P* < .05, ** *P* < .01, paired two-tailed student’s *t* test. MLN, mesenteric lymph nodes; siLP, small intestine lamina propria; cLP, colon lamina propria.

**Extended Data Figure 9. CAR expression and function in mucosal Teff cells is increased in response to intestinal inflammation. (a)** Percentages of CD3^+^CD4^+^ T cells in tissues of *Rag1*^−/−^ mice transplanted with congenic mixtures of wild type and CAR-deficient naïve CD4^+^ T cells over time, determined by flow cytometry. Representative of 5 mice per tissue and time point. **(b)** Mean absolute numbers of CD3^+^CD4^+^ T helper (T_H_) cells (± SEM; *n* = 5) in tissues of transferred B6.*Rag1*^−/−^ mice over time, determined by flow cytometry as in (a). **(c)** Mean relative *ex vivo* CAR (*Nr1i3*), MDR1 (*Abcb1a*), *Cyp2b10*, or *Il10* gene expression (± SEM; *n* = 3), determined by qPCR, in wild type (B6) CD4^+^ effector/memory (Teff) cells (sorted as in Extended Data Fig. 8a) from spleens of transferred B6.*Rag1*^−/−^ mice over time. **(d)** *Top row*, expression of Foxp3 and RORγt, determined by intracellular staining after *ex vivo* (PMA+ionomycin) stimulation, in CD4^+^CD44^hi^ cells from spleen (*left*) or small intestine lamina propria (siLP, *right*) of wild type (B6, blue) or CAR-deficient (B6.*Nr1i3*^−/−^, red) mice injected with isotype control (IgG) or soluble anti-CD3. *Bottom 4 rows*, expression of IL-10 and IL-17A in wild type or CAR-deficient spleen or siLP T cell subsets from mice treated +/− isotype control (IgG) or anti-CD3 antibodies. Cells were gated and analyzed by flow cytometry as above. Numbers indicate percentages; representative of 3 mice per group and genotype analyzed over 2-independent experiments. **(e-f)** Mean percentages of IL-10-expressing T cell subsets (± SEM; *n* = 3), gated and analyzed by *ex vivo* flow cytometry as in (a), in spleen **(f)** or siLP **(e)** T_H_ cell subsets from wild type (B6, blue) or CAR-deficient (B6.*Nr1i3*^−/−^, red) mice injected with or without isotype control (IgG) or anti-CD3 antibody. * *P* < .05, one-unpaired student’s *t* test; some *P* values are listed directly.

**Extended Data Figure 10. TCPOBOP protection against bile acid-induced ileitis requires CAR expression in T cells. (a)** Mean weight loss (± SEM; *n* = 5/group) of co-housed B6.*Rag2*^−/−^ mice transplanted with CAR-deficient (B6.*Nr1i3*^−/−^) CD4^+^ naïve T cells and maintained on a CA-supplemented diet with (CA/TC) or without (CA/Veh) TC treatment. Weights are shown relative to 3-weeks post-transfer when TC treatments were initiated. NS, not significant; two-way ANOVA. **(b)** H&E-stained sections of terminal ilea or colons from B6.*Rag2*^−/−^ mice reconstituted with CAR-deficient T cells and treated as above and as indicated. Representative of 5 mice/group. **(c)** Mean histology scores (± SEM) for colons or terminal ilea as in (b). NS, not significant; paired student’s *t* test.

## Notes

### Summary of Updates

This updated version contains additional information in main Figure 2d concerning genes and pathways activated downstream of CAR/Nr1i3 in small intestinal T cells. It also contains an additional supplementary table (2), which lists relevant gene sets from our statistical analyses of RNA-seq data in Figure 2a-d.

## References

1 Hofmann, A. F. & Hagey, L. R. Key discoveries in bile acid chemistry and biology and their clinical applications: history of the last eight decades. Journal of lipid research 55, 1553–1595, doi:10.1194/jlr.R049437 (2014).

2 Poupon, R., Chazouilleres, O. & Poupon, R. E. Chronic cholestatic diseases. J Hepatol 32, 129–140 (2000).

3 Arab, J. P., Karpen, S. J., Dawson, P. A., Arrese, M. & Trauner, M. Bile acids and nonalcoholic fatty liver disease: Molecular insights and therapeutic perspectives. Hepatology 65, 350–362, doi:10.1002/hep.28709 (2017).

4 Cao, W. et al. The Xenobiotic Transporter Mdr1 Enforces T Cell Homeostasis in the Presence of Intestinal Bile Acids. Immunity 47, 1182–1196 e1110, doi:10.1016/j.immuni.2017.11.012 (2017).

5 Lazar, M. A. Maturing of the nuclear receptor family. J Clin Invest 127, 1123–1125, doi:10.1172/JCI92949 (2017).

6 Ludescher, C. et al. Detection of activity of P-glycoprotein in human tumour samples using rhodamine 123. British journal of haematology 82, 161–168 (1992).

7 Pols, T. W., Noriega, L. G., Nomura, M., Auwerx, J. & Schoonjans, K. The bile acid membrane receptor TGR5 as an emerging target in metabolism and inflammation. J Hepatol 54, 1263–1272, doi:10.1016/j.jhep.2010.12.004 (2011).

8 Zhang, J., Huang, W., Qatanani, M., Evans, R. M. & Moore, D. D. The constitutive androstane receptor and pregnane X receptor function coordinately to prevent bile acid-induced hepatotoxicity. J Biol Chem 279, 49517–49522, doi:10.1074/jbc.M409041200 (2004).

9 Cerveny, L. et al. Valproic acid induces CYP3A4 and MDR1 gene expression by activation of constitutive androstane receptor and pregnane X receptor pathways. Drug Metab Dispos 35, 1032–1041, doi:10.1124/dmd.106.014456 (2007).

10 Wei, P., Zhang, J., Egan-Hafley, M., Liang, S. & Moore, D. D. The nuclear receptor CAR mediates specific xenobiotic induction of drug metabolism. Nature 407, 920–923, doi:10.1038/35038112 (2000).

11 Evans, R. M. & Mangelsdorf, D. J. Nuclear Receptors, RXR, and the Big Bang. Cell 157, 255–266, doi:10.1016/j.cell.2014.03.012 (2014).

12 Staudinger, J. L. et al. The nuclear receptor PXR is a lithocholic acid sensor that protects against liver toxicity. Proc Natl Acad Sci U S A 98, 3369–3374, doi:10.1073/pnas.051551698 (2001).

13 Ostanin, D. V. et al. T cell transfer model of chronic colitis: concepts, considerations, and tricks of the trade. American journal of physiology. Gastrointestinal and liver physiology 296, G135–146, doi:10.1152/ajpgi.90462.2008 (2009).

14 Arnold, M. A. et al. Colesevelam and Colestipol: Novel Medication Resins in the Gastrointestinal Tract. The American journal of surgical pathology, doi:10.1097/PAS.0000000000000260 (2014).

15 Dawson, P. A., Lan, T. & Rao, A. Bile acid transporters. Journal of lipid research 50, 2340–2357, doi:10.1194/jlr.R900012-JLR200 (2009).

16 Cui, J. Y. & Klaassen, C. D. RNA-Seq reveals common and unique PXR- and CAR-target gene signatures in the mouse liver transcriptome. Biochim Biophys Acta 1859, 1198–1217, doi:10.1016/j.bbagrm.2016.04.010 (2016).

17 Niu, B. et al. In vivo genome-wide binding interactions of mouse and human constitutive androstane receptors reveal novel gene targets. Nucleic Acids Res 46, 8385–8403, doi:10.1093/nar/gky692 (2018).

18 De Calisto, J. et al. T-cell homing to the gut mucosa: general concepts and methodological considerations. Methods Mol Biol 757, 411–434, doi:10.1007/978-1-61779-166-6_24 (2012).

19 Maglich, J. M. et al. Identification of a novel human constitutive androstane receptor (CAR) agonist and its use in the identification of CAR target genes. J Biol Chem 278, 17277–17283, doi:10.1074/jbc.M300138200 (2003).

20 Ramesh, R. et al. Pro-inflammatory human Th17 cells selectively express P-glycoprotein and are refractory to glucocorticoids. J Exp Med 211, 89–104, doi:10.1084/jem.20130301 (2014).

21 Moore, L. B. et al. Pregnane X receptor (PXR), constitutive androstane receptor (CAR), and benzoate X receptor (BXR) define three pharmacologically distinct classes of nuclear receptors. Mol Endocrinol 16, 977–986, doi:10.1210/mend.16.5.0828 (2002).

22 Karwacz, K. et al. Critical role of IRF1 and BATF in forming chromatin landscape during type 1 regulatory cell differentiation. Nat Immunol 18, 412–421, doi:10.1038/ni.3683 (2017).

23 Gagliani, N. et al. Coexpression of CD49b and LAG-3 identifies human and mouse T regulatory type 1 cells. Nat Med 19, 739–746, doi:10.1038/nm.3179 (2013).

24 Korn, T., Bettelli, E., Oukka, M. & Kuchroo, V. K. IL-17 and Th17 Cells. Annu Rev Immunol 27, 485–517, doi:10.1146/annurev.immunol.021908.132710 (2009).

25 Maynard, C. L. et al. Regulatory T cells expressing interleukin 10 develop from Foxp3+ and Foxp3-precursor cells in the absence of interleukin 10. Nat Immunol 8, 931–941, doi:10.1038/ni1504 (2007).

26 Sano, T. et al. An IL-23R/IL-22 Circuit Regulates Epithelial Serum Amyloid A to Promote Local Effector Th17 Responses. Cell 163, 381–393, doi:10.1016/j.cell.2015.08.061 (2015).

27 Wan, Q. et al. Cytokine signals through PI-3 kinase pathway modulate Th17 cytokine production by CCR6+ human memory T cells. J Exp Med 208, 1875–1887, doi:10.1084/jem.20102516 (2011).

28 Yadava, K. et al. Natural Tr1-like cells do not confer long-term tolerogenic memory. Elife 8, doi:10.7554/eLife.44821 (2019).

29 Barrat, F. J. et al. In vitro generation of interleukin 10-producing regulatory CD4(+) T cells is induced by immunosuppressive drugs and inhibited by T helper type 1 (Th1)- and Th2-inducing cytokines. J Exp Med 195, 603–616, doi:10.1084/jem.20011629 (2002).

30 Song, X. et al. Microbial bile acid metabolites modulate gut RORgamma(+) regulatory T cell homeostasis. Nature 577, 410–415, doi:10.1038/s41586-019-1865-0 (2020).

## References

31 Kim, J. J., Shajib, M. S., Manocha, M. M. & Khan, W. I. Investigating intestinal inflammation in DSS-induced model of IBD. J Vis Exp, doi:10.3791/3678 (2012).

32 Berg, D. J. et al. Enterocolitis and colon cancer in interleukin-10-deficient mice are associated with aberrant cytokine production and CD4(+) TH1-like responses. J Clin Invest 98, 1010–1020, doi:10.1172/JCI118861 (1996).

33 Langmead, B. & Salzberg, S. L. Fast gapped-read alignment with Bowtie 2. Nature methods 9, 357–359, doi:10.1038/nmeth.1923 (2012).

34 Zhang, Y. et al. Model-based analysis of ChIP-Seq (MACS). Genome Biol 9, R137, doi:10.1186/gb-2008-9-9-r137 (2008).

35 Robinson, J. T. et al. Integrative genomics viewer. Nat Biotechnol 29, 24–26, doi:10.1038/nbt.1754 (2011).

36 Durant, L. et al. Diverse targets of the transcription factor STAT3 contribute to T cell pathogenicity and homeostasis. Immunity 32, 605–615, doi:10.1016/j.immuni.2010.05.003 (2010).

37 Ritchie, M. E. et al. limma powers differential expression analyses for RNA-sequencing and microarray studies. Nucleic Acids Res 43, e47, doi:10.1093/nar/gkv007 (2015).

38 Wei, G. et al. Global mapping of H3K4me3 and H3K27me3 reveals specificity and plasticity in lineage fate determination of differentiating CD4+ T cells. Immunity 30, 155–167, doi:10.1016/j.immuni.2008.12.009 (2009).

39 Yusuf, I. et al. Germinal center T follicular helper cell IL-4 production is dependent on signaling lymphocytic activation molecule receptor (CD150). Journal of immunology 185, 190–202, doi:10.4049/jimmunol.0903505 (2010).

40 Suino, K. et al. The nuclear xenobiotic receptor CAR: structural determinants of constitutive activation and heterodimerization. Mol Cell 16, 893–905, doi:10.1016/j.molcel.2004.11.036 (2004).

